# Vascular dysfunction in Huntington’s disease is located at the blood-CSF barrier and is rescued by sphingosine-1-phosphate receptor agonist

**DOI:** 10.1101/2025.04.30.650996

**Authors:** Krzysztof Kucharz, Rana Soylu-Kucharz, Martine Saint-Pierre, Melanie Alpaugh, Filippa L. Qvist, Maria Björkqvist, Niels Henning Skotte, Francesca Cicchetti, Martin Lauritzen

## Abstract

Brain vascular barriers are dysfunctional in many neurological disorders, including Huntington’s disease (HD), but which vessels are affected and how remains unclear. Using in vivo two-photon microscopy in R6/2 HD-model mice, we reveal a striking divergence in barrier dysfunction. Surface vessels of the blood-cerebrospinal fluid barrier (BCSFB) drive pathology by paracellular leakage, while the blood-brain barrier (BBB) of parenchymal arterioles and capillaries exhibited increased vesicular transport and adsorptive-mediated transcytosis (AMT), with arterioles being the most susceptible segment. The pathology of both barriers was congruent with astrocyte activation localized to affected vessels in R6/2 mice, as well as in post-mortem HD human brains. Sphingosine-1-phosphate receptor 1 (S1PR1) agonist treatment rescued BCSFB leakage but only partially restored BBB function, selectively reducing AMT in arterioles, with no effect in capillaries and venules. These findings redefine HD vascular pathology, demonstrating heterogeneous, vessel-type-specific barrier failure beyond generalized “disruption,” uncovering mechanistic vulnerabilities and the therapeutic potential of S1PR1 modulation.

## INTRODUCTION

In the human brain, surface (pial) vessels form the blood-cerebrospinal fluid barrier (BCSFB), whereas cerebral (parenchymal) vessels, primarily capillaries, constitute the blood-brain barrier (BBB). Capillaries directly contact brain tissue^1,2^, while the intermediate segments — penetrating arterioles and ascending venules — exhibit features of both BCSFB and BBB, at least at the level of transcriptomics^3–5^. Although capillaries are widely used in vitro to model vascular transport, this approach is unfit to reflect the functional heterogeneity of the microvascular network. For example, non-specific vesicular transport of proteins, adsorptive-mediated transcytosis (AMT), is heavily suppressed in capillaries but active in arterioles^6,7^, while immune cell trafficking and receptor-mediated transcytosis (RMT) of macromolecules occur primarily in venules^2,8^.

In pathology, dysfunction of the BCSFB/BBB is often described as the disruption of junctional complexes. This view oversimplifies that distinct vessel types serve specialized transport roles^7–10^. Thus, changes in barrier permeability may proceed via different mechanisms depending on vessel type, a critical nuance that remains underexplored despite its clear implications for diagnostic accuracy and therapeutic targeting in human disease contexts. Here, we address this gap by characterizing vessel-type-specific barrier pathologies in Huntington’s disease (HD), a fatal neurodegenerative disorder. with urgent clinical need for disease-modifying interventions.

HD leads to the toxic accumulation of mutant huntingtin (mHtt)^11^, triggering progressive neuronal dysfunction and brain atrophy^12–15^. The mHtt is expressed throughout the microvascular network — including arterioles, capillaries, and venules—where it forms aggregates in brain endothelial cells (BECs), the principal component of the BBB and BCSFB, as well as in pericytes, astrocytes, and smooth muscle cells^16–20^. These microvascular abnormalities manifest early, preceding the onset of motor, cognitive, and metabolic symptoms in HD^21,22^, making them a potentially valuable target for early intervention.

Using in vivo two-photon microscopy (2PM) in R6/2 transgenic HD mice and human post-mortem tissue analyses, we provide a detailed account of BCSFB and BBB dysfunction, offering the first insights into vessel-type-specific alterations in paracellular leakage and vesicular transport in HD. We demonstrate that in HD mice, barrier dysfunction follows a distinct pattern: BCSFB paracellular leakage is confined to large subpial vessels, while the paracellular barrier is preserved at capillaries and parenchymal arterioles. In comparison, vesicular transport by AMT increases up to 8-fold in parenchymal arterioles and 2.2-fold in brain capillaries. Treatment with an S1PR1 agonist rescued BCSFB integrity and reduced AMT in arterioles but failed to restore BBB function in capillaries and venules. Astrocyte activation matched the topology of barrier breakdown, both in mice and brains of HD patients, clustering around hot-spots of barrier leakage, with the most pronounced effects in female patients. These findings suggest that brain barrier defects in HD patients are a treatable condition, holding promise for the development of more effective therapeutics for HD.

## RESULTS

### BCSFB drives paracellular leakage in R6/2 mice

BBB dysfunction manifests as increased paracellular permeability between adjoining brain endothelial cells (BECs), allowing blood-borne substances to invade the brain. To spatially map leakage into the brain parenchyma, we used two-photon fluorescence microscopy (2PM) *in vivo* in R6/2 mice (Fig. 1A). We intravenously injected Sodium Fluorescein (NaFluo, 0.376 kDa), a sensitive hydrophilic fluorophore for 2PM barrier permeability assays^6,23,24^ (Fig. 1B). We collected hyperstack (Z-stack over time) data from anatomically distinct cortical regions^25^: the sub-pial volume (0-60 µm), primarily containing large vessels forming the BCSFB, and the capillary bed volume (120-180 µm) with vessels forming the BBB (Fig. 1C). R6/2 mice exhibited increased paracellular leakage at the sub-pial volume, indicated by substantial NaFluo accumulation, suggesting BCSFB dysfunction (Fig. 1D,E; Supplementary Video 1). In contrast, paracellular leakage at the capillary bed remained unchanged (Fig. 1D,F; Supplementary Video 1). To calculate permeability, we extracted NaFluo signals from blood vessels over time (Fig. 1G, Extended Data Fig. 1), calculated the average AUC of the brain/blood ratio (Fig. 1H,I)^24^. Our results showed a ∼3.8-fold increase in sub-pial paracellular permeability in R6/2 mice compared to WT, with no significant change at the capillary BBB (Fig. 1H,I; Supplementary Table 1A,B). Thus, the BCSF barrier-containing vessels, rather than BBB-containing capillaries, drive the pathological increase in paracellular permeability.

**Figure 1.**
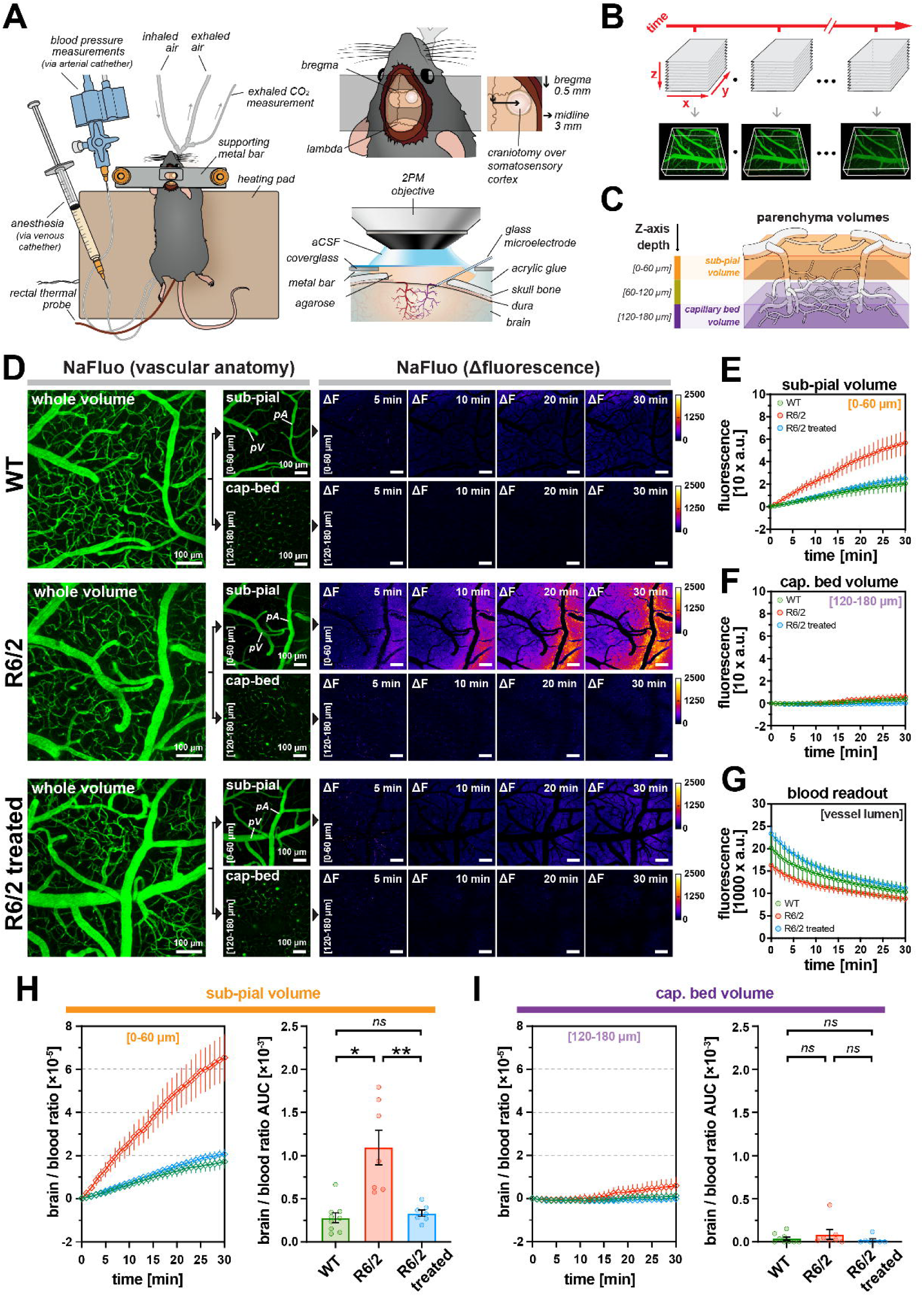
Pial vessels drive vascular leakage pathology in R6/2 HD mice. **(A)** Schematic drawing of a mouse following the microsurgery for 2-photon imaging in vivo. **(B)** The principle of hyperstack imaging. The fluorescence signal is recorded as Z-stack images over time, i.e., four-dimensional volumetric data (xyzt). The microvasculature is delineated by i.v.-injected blood-circulating NaFluo (green). **(C)** Schematic representation of the imaged cortical parenchyma. The brain volume was divided into *sub-pial* volume (0-60 µm below pia) occupied mainly by large pial vessels and sinuses forming BCSF, and *capillary bed* volume (120-180µm below pia) occupied mainly by capillaries forming BBB. **(D)** Left large panels: maximum-intensity projected Z-stacks showing anatomical features of the imaged brain area. The vessels are delineated by i.v.-injected blood-circulating NaFluo (1% 40 µL /20 ganimal). Small panels: average-intensity projected Z-stacks from *sub-pial* and *capillary bed* volumes (NaFluo signal in green) and their respective timelapse data showing gradual accumulation of NaFluo in the brain parenchyma, measured as fluorescence intensity increase (ΔF, arbitrary fluorescence units, a.u.) after subtraction of the baseline (t=0 min) and represented pseudo-color scale (ImageJ, fire LUT). Compared to WT animals, the R6/2 mice exhibit increased paracellular leakage of blood-circulating NaFluo, with the effect present only in *sub-pial* volume. Treatment of R6/2 mice with S1PR1 agonist SEW2981 reverses the pathological leakage across the BCSF. **(E)** Quantitative assessment of NaFluo increase in the brain parenchyma *sub-pial* volume and **(F)** *capillary bed* volume, **(G)** with concurrent measurement of decrease of NaFluo circulating in the bloodstream. **(H-I)** Quantification of paracellular leakage of NaFluo into the brain in *sub-pial* and *capillary bed* volumes. The brain/blood ratio traces show NaFluo signal intensity increase after normalization of NaFluo signal increase from panel E) for *sub-pial,* and panel F) for *capillary bed* volume to AUC of NaFluo signal intensity in the blood from panel G). The bar graphs show the AUC calculated from the obtained brain/blood ratio traces. **<u>All panels</u>**: nWT= 9; nR6/2= 7; nR6/2+= 7 mice; data is shown as average ± SEM; two-tailed t-test; *=p<0.05; **=*p*<0.01; *ns*=non-significant; *aCSF* = artificial cerebrospinal fluid, *NaFluo* = Sodium Fluorescein, *LUT =* look-up table, *a.u.* = arbitrary fluorescence units (detector counts), *cap-bed* = capillary bed volume, *AUC* = area under curve.

### Stimulating S1PR1 restores normal BBB permeability

Sphingosine-1-phosphate (S1P) is the blood-borne signalling molecule activating S1P receptor 1 (S1PR1) at the luminal side of BECs^26^. Ongoing activity of S1PR1 maintains low paracellular permeability of endothelial barriers, making S1PR1 signalling a targetable pathway in CNS vascular regulation, as evidenced in mice with BBB deficits^6,26,27^, and with the clinical success of the S1PR1 agonist fingolimod in treating ALS^28^. Here, we investigated whether S1PR1 agonism reversed BCSFB pathology in HD mice^6^. Mice received intraperitoneal SEW2871 (10 μg/g) for six days at 24-hour intervals, with the final dose on the imaging day. The treatment restored the low BCSFB permeability in R6/2 mice, reducing it to levels of WT, healthy mice (Fig. 1D-F; Supplementary Table 1C,D). Thus, impairment of the S1PR1 pathway contributes to endothelial dysfunction in the R6/2 mice model of HD, which can be mitigated by S1PR1 agonism.

### Arterioles are most susceptible to AMT increase

AMT transports plasma proteins, such as albumin, across the BBB^10,29^. In healthy brains, AMT is suppressed by S1P in blood and the fatty acid transporter mfsd2a in brain endothelial cells^6,30,31^. Disinhibition of AMT is an early marker of BBB dysfunction preceding ultrastructural alterations and paracellular leaks, e.g., in aging^10^ and stroke^32^. Although an increased presence of vesicular structures was detected in post-mortem HD brains^16^, its link to AMT remained unclear. Using 2PM, we mapped AMT in vivo in R6/2 mice across distinct vessel types. Mice were injected with fluorescent bovine serum albumin (BSA-A488), and its uptake in BECs was tracked over time (Fig. 2A). Within 120 min post-injection, we observed a gradual increase in AMT vesicles filled with BSA-A488 endocytosed from the bloodstream (Fig. 2A, Supplementary Video 2; Extended Data Fig. 2A, B). Notably, we observed a gradual uptake of transcytosed albumin in cells (Supplementary Video 3), identified as perivascular macrophages^6^. The subsequent 70kDa TRITC-dx injection outlined the vessel lumen, allowing for clear differentiation of AMT vesicles from autofluorescent debris (Fig. 2B,C). In WT mice, AMT was uniformly low across all vessel types. In contrast, R6/2 mice showed widespread AMT disinhibition, with an overall ∼3-fold increase (Fig. 2D,E; Supplementary Table 1E). Specifically, penetrating arterioles (*penA*) were the most affected, exhibiting an ∼8-fold AMT increase, pial arterioles, pial venules, and post-capillary/ascending venules had a moderate ∼4-fold increase, and capillaries and sinuses were least affected, with a relatively modest ∼2-fold increase (Fig. 2F, Extended Data Fig. 2B; Supplementary Table 1F-K). Notably, this increase was not due to, e.g., blood-flow stalling, but reflected a true rise in vesicular transport, as stalled capillaries did not show puncta enrichment (Extended Data Fig. 2D). Thus, pathologic AMT disinhibition occurs across all vessel types in HD, affecting both the BCSFB and BBB. However, the extent of dysfunction varies, with penetrating arterioles being the most vulnerable and capillaries the least affected, highlighting vessel-specific heterogeneity in BBB transport pathology.

**Figure 2.**
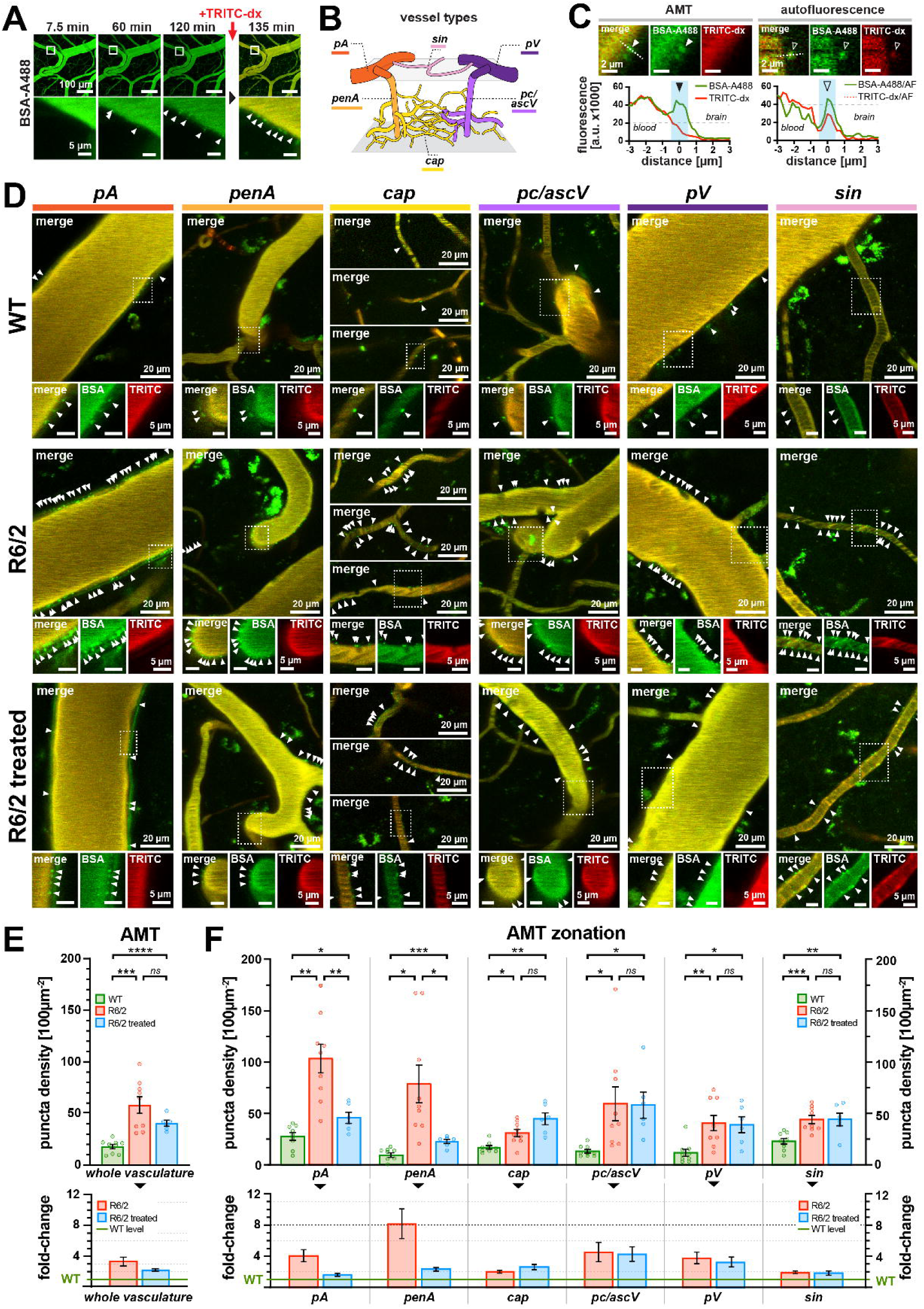
Vascular zonation of AMT pathology, and heterogenous rescue potential of BCSF and BBB in R6/2 HD mice. **(A)** Timelapse images of the gradual uptake of blood-circulating BSA-A488 (green) into AMT vesicular punctae at the BCSF/BBB interface. The arrowheads (white) in magnified panels show the individual AMT vesicular punctae (green). The vessel lumen is delineated by injecting TRITC-dx (red) 120 minutes after BSA-A488. **(B)** Anatomical representation of the division of the cortical blood microvasculature into major vessel types: pial arterioles (*pA*), penetrating arterioles (*penA)*, capillaries *(cap)*; post-capillary/ascending venules *(pc/ascV)*, pial venules (*pV*), and sinuses *(sin)*. **(C)** The distinction between AMT vesicular punctae and non-specific brain autofluorescence. The AMT punctae (left panel) exhibit only the signal from BSA-A488 (green), whereas the autofluorescence signal (AF) also contains the elements of non-specific signal in the red emission range (red). The dashed line shows the axis of the fluorescence intensity profiles presented in underneath panels. The filled arrowhead shows the location AMT puncta, the empty arrowhead shows ‘false’ punctae not classified as AMT. **(D)** Detailed images showing AMT punctae (arrowheads) at distinct vessel types, outlined in panel B. The images were collected 135 min post-BSA-A488 injection, with TRITC-dx delineating the vessel lumen. WT animals show a low degree of AMT across all vessel types, in contrast to R6/2 mice exhibiting overt pathological disinhibition of AMT, with the limited effect of the S1PR1-targeted treatment in pial and penetrating arterioles. **(E-F)** The upper panel shows the absolute AMT count, the lower panel shows a AMT levels fold change in relation to the WT baseline (=1). **(E)** Quantitative analyses of AMT puncta density (amount of punctae / vessel surface) showing the general increase in AMT in R6/2 mice and the general effect of the S1PR1-targeted treatment in cortical vasculature. **F)** Detailed, vessel-type-dependent analysis of AMT reveals the zonation vascular susceptibility to AMT increase in R6/2 mice and heterogeneous response to the treatment. The most sensitive are penetrating arterioles (*penA)*, the least sensitive are sinuses *(sin)* and capillaries *(cap)*. **<u>All panels</u>**: nWT= 9; nR6/2= 9; nR6/2+= 6 mice; data is shown as average ± SEM; two-tailed t-test; *=p<0.05; **=*p*<0.01; ***=*p*<0.001; ****=p<0.0001; *ns*=non-significant.

### S1PR1 agonism suppresses AMT in arterioles but not in other vessel types

In addition to regulating paracellular permeability, S1P–S1PR1 signalling agonists have been shown to suppress AMT in inflammatory-like conditions^6^. Therefore, we next investigated whether S1PR1 agonist treatment alleviated the increase in AMT in the HD mice. We used the same mice as in the paracellular permeability assay, i.e., treated over six days with SEW2871 (10 μg/g body weight). Treated R6/2 mice showed an overall reduction in endothelial AMT (Fig. 2D,E). However, the effect was restricted to pial and penetrating arterioles. In these vessels, AMT levels were significantly decreased (Fig. 2F; Supplementary Table 1L, M), from ∼4 to ∼1.7-fold in pial arterioles and from ∼8-fold to 2.4-fold in penetrating arterioles compared to WT baseline (Fig. 2F; Supplementary Table 1N, O). In contrast, the treatment was ineffective for capillaries, post-capillary/ascending venules, pial venules, and sinuses (Fig. 2F, Supplementary Table 1P-S). Thus, the rise in AMT is heterogeneous across cerebral microvessels in the HD model mouse, with important differences in rescue potential for the S1P-dependent mechanism for different vessel types.

### An inverse relation between paracellular permeability and AMT

In stroke, BBB dysfunction follows a biphasic pattern, where either AMT increases or paracellular leakage becomes more prominent at different stages^32^. It is unknown whether a similar pattern exists for chronic brain disorders, including HD. We cross-correlated paracellular leakage and AMT levels across cortical volumes in the same R6/2 mice, analyzing distinct vessel types within matched regions. We found an inverse correlation between paracellular leakage and AMT (Extended Data Fig. 3A). This was evident in pial arterioles within the subpial volume, where animals with high paracellular leakage exhibited only a moderate increase in AMT and vice versa, but not in less affected vessels, i.e., pial venules and sinuses that occupy the same parenchymal niche and capillaries (Extended Data Fig. 3B-D). Thus, in the most vulnerable vessels, arterioles exhibit a reciprocal pattern where either AMT or paracellular permeability predominates, suggesting a dynamic interplay between two distinct mechanisms of BBB dysfunction.

### Proteome signature highlights heterogeneous BBB/BCSFB and vessel pathology

Next, we aimed to explore the molecular basis of the increase in paracellular and transcellular transport in the HD mice. For this purpose, we used mass spectrometry to analyse blood plasma and CSF samples collected from mice treated over six days with SEW2871 (10 μg/g body weight)(Fig. 1A). The obtained profiles of plasma and CSF revealed heterogeneous protein signatures, implicating dysfunction of both the BBB and BCSFB across arterioles, capillaries, and venules (Fig. 3B-E, Extended Data Fig. 4). Unsupervised clustering distinguished WT, R6/2, and treated R6/2 groups, suggesting broad yet distinct proteomic shifts across barrier compartments. Elevated CXCL12, CD68, and C1q in R6/2 CSF indicated immune infiltration and complement-driven endothelial injury, particularly at venules, consistent with paracellular leakage (Table 1). Simultaneously, increased Rab8b, SUMF1/2, and VEGFC supported the active involvement of adsorptive-mediated transcytosis (AMT), particularly in arterioles, suggesting vesicle-driven transport defects in HD. This dual mechanism—junctional disruption and vesicular overactivity— appeared to be spatially segregated, with AMT enrichment in arterioles and leukocyte-associated leakage in venules. Treatment with an S1PR1 agonist partially normalized this pattern (Fig. 3B-E), Rab8b and Plaur levels decreased, indicating reduced vesicle trafficking and ECM degradation, while CXCL10 and CD14 suppression reflected tighter junctional integrity and limited immune cell access (Table 2). Altogether, these mechanistic findings corroborate 2PM data, supporting the idea that barriers dysfunction is not uniform but varies by vessel type, vascular segment, and mechanism in HD.

**Figure 3.**
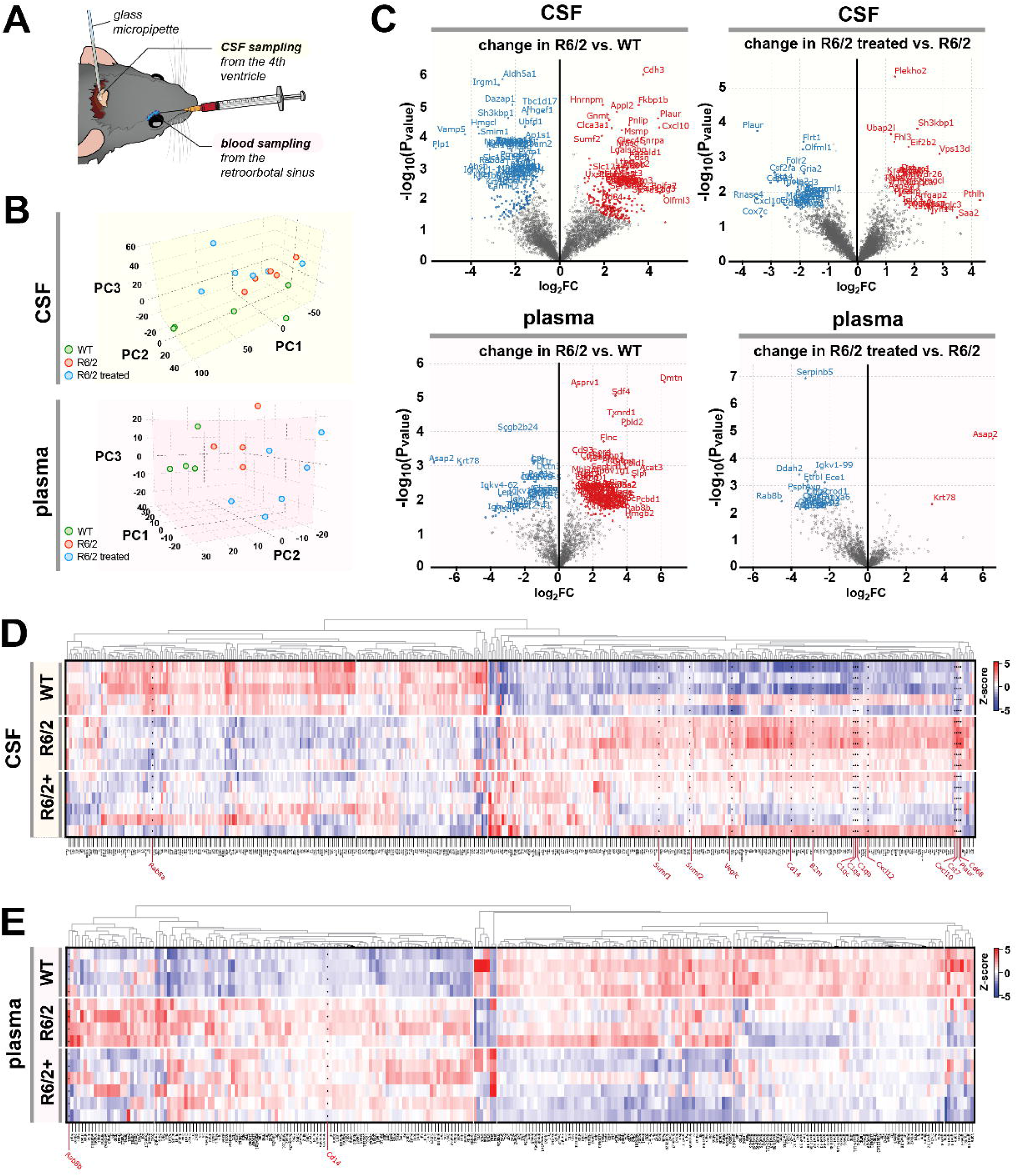
Blood plasma and CSF mass spectrometry fingerprinting reveal distinct clustering of proteins relevant for barriers pathology in HD. (A) Schematic drawing of a mouse head illustrating locations of CSF and blood collection for mass spectrometry analyses (**B**) PCA (principal-component analysis) demonstrating the difference between WT and R6/2 as well as the effect of S1PR1 agonist treatment measured in both plasma and CSF. **(C)** The volcano plots illustrate the significant protein expression changes between the two genotypes for both plasma and CSF, as well as the induced changes by treatment in both biofluids (FDR > 0.05; blue and red colors illustrate decreased and increased protein expression, respectively). **(D-E)** CSF and plasma cluster analysis based on significant hits demonstrating the grouping of the replicate samples. Highlighted proteins are of direct relevance to BCSFB/BBB function. nWT= 5; nR6/2= 5; nR6/2+= 6 mice. Two mice were excluded from plasma analysis due to low protein yield and protein identifications.

**Table 1:**
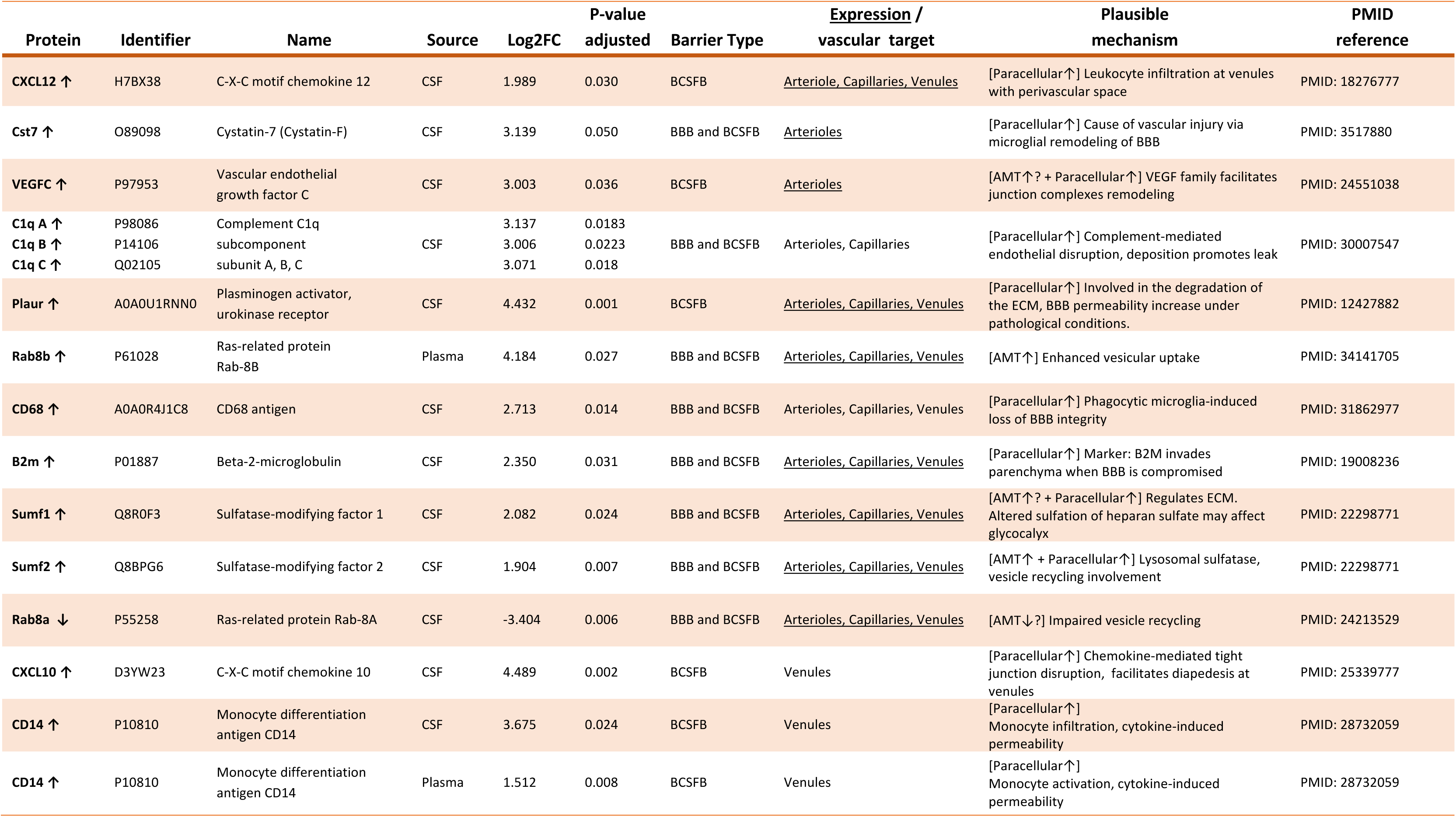
Proteome signature of R6/2 mice. MS fingerprint of selected blood/CSF proteins significantly elevated or decreased in HD (R6/2 vs. WT) relevant for BBB/BCSFB function. The suggested affected vessel types are based on the preferential protein expression in particular vessel types and/or converging mechanisms from the literature (PMID references). *<u>Underline</u>* indicates transcriptome expression in endothelial cells at respective vascular segments (PMID: 29443965). *ECM* = extracellular matrix, *TJ* = tight junctions, *AMT* = adsorptive-mediated transcytosis.

**Table 2:**
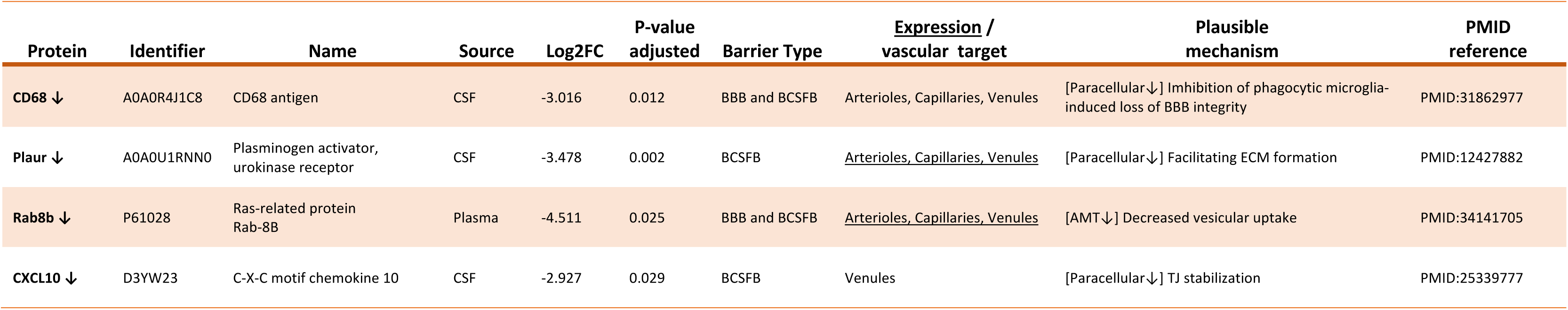
Proteome signature of S1PR1-agonist treated HD mice with implications for barrier restoration or further modulation. . MS fingerprint of selected proteins significantly elevated or decreased in blood or CSF of R6/2 treated vs. R6/2 mice relevant for BBB/BCSFB function. The suggested affected vessel types are based on the preferential protein expression in particular vessel types and/or converging mechanisms from the literature (PMID sources). *<u>Underline</u>* indicates transcriptome expression in endothelial cells at respective vascular segments (PMID: 29443965). *ECM* = extracellular matrix, *TJ* = tight junctions, *AMT* = adsorptive-mediated transcytosis.

| Protein | Identifier | Name | Source | Log2FC | P-value<br>adjusted | Barrier Type | <u>Expression /</u><br>vascular target | Plausible<br>mechanism | PMID<br>reference |
| --- | --- | --- | --- | --- | --- | --- | --- | --- | --- |
| CD68 ↓ | A0A0R4J1C8 | CD68 antigen | CSF | -3.016 | 0.012 | BBB and BCSFB | Arterioles, Capillaries, Venules | [Paracellular↓] Imhibition of phagocytic microglia-induced loss of BBB integrity | PMID:31862977 |
| Plaur ↓ | A0A0U1RNN0 | Plasminogen activator, urokinase receptor | CSF | -3.478 | 0.002 | BCSFB | <u>Arterioles, Capillaries, Venules</u> | [Paracellular↓] Facilitating ECM formation | PMID:12427882 |
| Rab8b ↓ | P61028 | Ras-related protein Rab-8B | Plasma | -4.511 | 0.025 | BBB and BCSFB | <u>Arterioles, Capillaries, Venules</u> | [AMT↓] Decreased vesicular uptake | PMID:34141705 |
| CXCL10 ↓ | D3YW23 | C-X-C motif chemokine 10 | CSF | -2.927 | 0.029 | BCSFB | Venules | [Paracellular↓] TJ stabilization | PMID:25339777 |

### Astrocyte activation mirrors patterns of BBB dysfunction

Next, we examined whether localized BCSFB/BBB disruption leads to spatially confined pathological responses in the adjacent brain parenchyma, specifically by activating astrocytes at sites of barrier compromise. Astrocytes react to BBB disruption by increasing glial fibrillary acidic protein (GFAP) expression, making them a reliable marker of vascular abnormalities^33^. Using immunohistochemistry, we visualized cerebral blood vessels (CD31), differentiated arterioles from venules (α-SMA), and assessed GFAP+ astrocyte activation in WT and R6/2 brain sections (Fig. 4A). GFAP+ astrocytes were increased in R6/2 mice, aligning with regions of paracellular leakage detected via 2PM (Fig. 1). To quantify this response, we stratified cortical regions into 60 µm increments based on exposure to leakage from the BCSFB and BBB in 2PM experiments, enabling a direct comparison of astrocyte activation across these zones (Fig1, Fig. 4B, Extended Data Fig.5).

**Figure 4.**
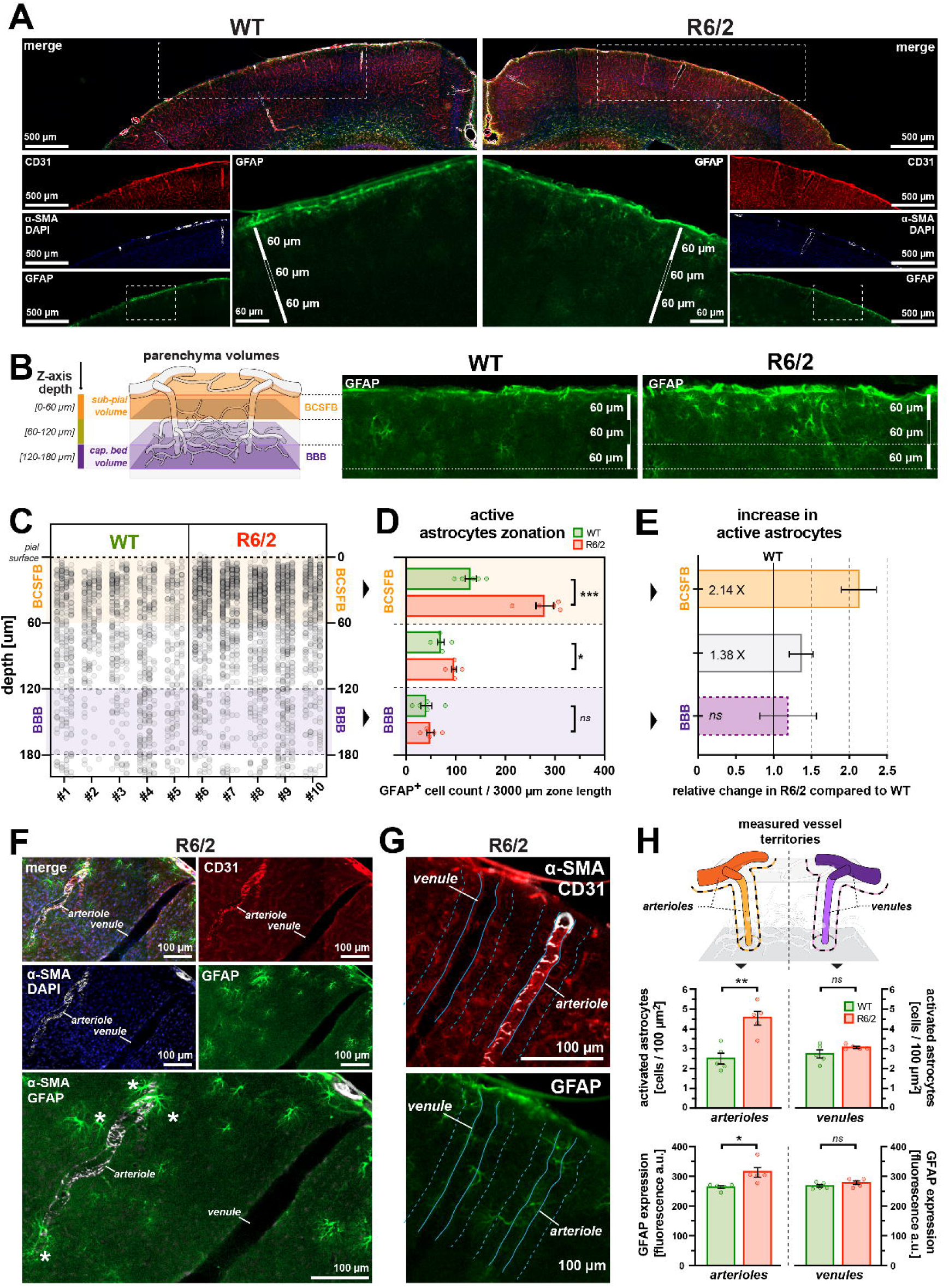
Location of activated astrocytes matches vascular topology of BBB disfunction. **(A)** Examples of immunostained sections showing cortical areas in WT and R6/2 mice. CD31 staining delineates vasculature (red), DAPI labels nuclei (blue), α-SMC labels arterioles (white), and GFAP stains activated astrocytes (green). **(B)** Distribution of GFAP+ astrocytes along a 3000 µm cortical segment. Each point represents an individual cell at its respective depth. Data from each mouse (#) include two sections (four hemispheres per mouse). **(D)** GFAP+ astrocytes show a preferential increase in R6/2 mice in the superficial cortical area (0-60 µm, BCSF influence), with no increase in numbers at the deeper layer (120-180 µm, capillary bed, BBB influence). **(E)** The Relative increase in GFAP+ astrocytes from panel (D) shows strong enrichment in superficial volumes, with no significant enrichment at the capillary bed volumes in R6/2 mice. **(F)** Example immunostaining images showing identification of arterioles by the presence of α-SMA signal, and accompanying GFAP+ astrocytes in proximity to vessels (asterisks) **(G)** Example of a cut-off point distance (20 µm) of measured vessel territories for arterioles and venules. **(H)** Upper panel: schematic image showing the measurement areas for arterioles and venules. Lower panels: in R6/2 mice, the GFAP+ astrocytes are present preferentially in proximity to arterioles, with a trend to enrichment in venules. Likewise, their GFAP expression levels measured by fluorescence signal intensity is increased at arterioles compared to WT, with no difference in venules. **<u>All panels:</u>** nWT= 5; nR6/2= 5 mice. Data is shown as average ± SEM; two-tailed t-test; *=p<0.05; **=*p*<0.01; ***=*p*<0.001; *ns*=non-significant.

Astrocyte activation followed a zonation effect, with GFAP+ astrocytes primarily enriched in cortical areas influenced by BCSFB leakage (0–60 µm). R6/2 mice exhibited a ∼2.14-fold increase in GFAP+ astrocytes compared to WT in these superficial cortical areas (Fig. 4C-E). In contrast, deeper regions associated with intact BBB function (120-180 µm, capillary bed) showed no significant astrocyte activation, suggesting that astrocyte responses are congruent with the affected barrier compartment (Fig. 4C-E). To further explore the spatial relationship between astrocyte activation and vascular dysfunction, we analyzed whether astrocytes were preferentially enriched near arterioles, the vessel type most susceptible to AMT disinhibition (Fig. 2). A standardized cutoff distance of 20 µm from the vessel wall was applied (Figure 4G), approximately twice the average estimated distance between astrocyte and its nearest vessel in the brain^25^. We observed a twofold increase in astrocyte density around arterioles in R6/2 mice (Fig. 4H), which contrasted with no significant changes around venules, reinforcing the link between astrocyte activation and barrier-compromised vessel types. Notably, we also detected elevated GFAP expression intensity, further confirming increased activation at sites of BBB dysfunction (Fig. 4H). To rule out a generalized increase in glial cell density as an explanation, we used 2PM and intracortical SR101 injections to measure in vivo astrocyte density. While R6/2 mice showed a modest ∼25% increase in total astrocyte density (Extended Data Fig. 6), this was insufficient to account for the twofold increase in GFAP+ astrocytes detected *ex vivo*, confirming that the observed enrichment results from localized barrier dysfunction rather than global glial changes in HD. Thus, the presence of astrocyte activation mirrors the vascular topology of the BBB dysfunction.

### Functional impact of BBB leakage in HD

Astrocytes actively participate in synaptic transmission, neuronal modulation, and neurovascular responses. Their activation has been linked to cortical hyperexcitability that exacerbates neurodegeneration in chronic and acute disorders, both in humans and mice^34–36^. Given that increased paracellular leakage coincided with GFAP+ astrocyte activation in superficial cortical layers (∼60 µm from the pia), we hypothesized that R6/2 mice may exhibit increased excitatory signalling and impaired inhibition, leading to hyperexcitability. To test this, we conducted electrical stimulation of the whisker pad, activating the somatosensory (barrel) cortex, where thalamic inputs project to superficial cortical layers—the site of astrocyte activation^37^.

Each mouse underwent three stimulation series across multiple frequencies while recording electrocorticogram (ECoG), local field potentials (LFPs), and direct current (DC) potentials from electrodes placed 50-70 µm into the barrel cortex. Additionally, we monitored mean arterial blood pressure (MABP) and exhaled CO₂ (exCO₂) as systemic controls (Fig. 5A-C). For each stimulation train, we extracted excitatory post-synaptic potential (EPSP) peak values and latencies and inhibitory post-synaptic potential (IPSP) peak values and latencies at frequencies 0.5 - 5 Hz (Fig. 5D,E). To assess cumulative excitatory and inhibitory activity, we summed the EPSPs and IPSPs (ΣEPSPs, ΣIPSPs) over the full 15-second stimulation duration (Fig. 5F). The results revealed that HD pathology leads to increased neuronal excitability, with a ∼2-fold increase in Σ_EPSPs_ across all stimulation frequencies (Fig. 5F, Supplementary Table 2A-B). Additionally, R6/2 mice displayed altered activation and inhibition latencies — EPSPs occurred earlier, while IPSPs were delayed (Fig. 5E,G). This faster excitatory activation and delayed inhibition resulted in an extended activation period, with a longer EPSP-IPSP interval in R6/2 mice (Fig. 5G, Supplementary Table 2D-F). Notably, we have observed instances of abnormal, spontaneous constrictions of pial arterioles in R6/2 mice, even without induced stimuli (Supplementary Video 4), which were absent in WT mice. Thus, the R6/2 mice’s thalamic-cortical route exhibited increased neuronal excitability, consistent with BCSFB dysfunction and localized glial activation in the superficial layers of the cortex, which are integral to this neurocircuitry.

**Figure 5.**
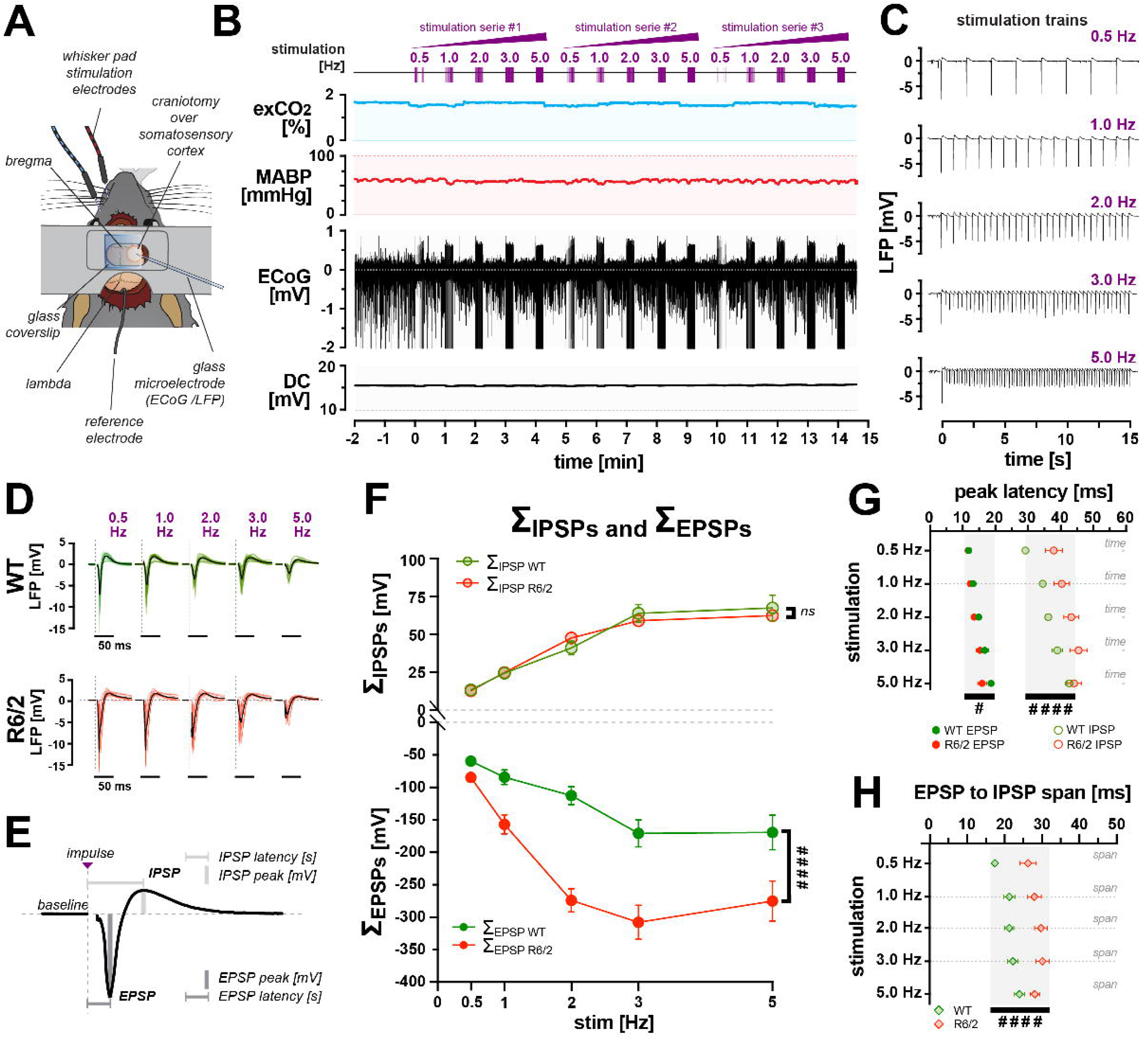
Hyperexcitability at the apical layers of the barrel cortex in R6/2 mice. **(A)** Schematic drawing showing the placement of electrodes for somatosensory stimulation of whisker barrel cortex. **(B)** Examples of concurrent recording of brain electrophysiological activity (electrocorticogram, *ECoG*), direct current *(DC)*, and accompanying systemic parameters, i.e., exhaled CO2 *(exCO2)*and mean arterial blood pressure *(MABP)*. The upper timeline shows the events of three series of 15 s stimulation trains ranging from 0.5 to 5.0 Hz. **(C)** Changes in local field potential *(LFP)* during trains of stimulations. **(D)** The mean traces of LFP responses for WT and R6/2 mice (black), with colored tracers representing individual mice. **(E)** The principal components of LFP response to somatosensory stimulation. Stimulation onset (*impulse)* triggers excitatory post-synaptic potential (*EPSP*), followed by compensating inhibitory post-synaptic potential (*IPSP).* Both EPSP and IPSP were measured for peak (degree of potential change) and peak latency (peak time from stimulation onset). **(F)** Quantitative analyses of the total sum of EPSPs and IPSPs for each stimulation frequency in WT and R6/2 mice. Compared to WT, the R6/2 mice show a significant increase in neuronal excitability (EPSPs) across all stimulation frequencies. **(G-H)** Temporal profiles of synaptic activation showing that R6/2 mice have a quicker initiation of excitatory postsynaptic potentials (EPSPs) and a pronounced lag in inhibitory postsynaptic potentials (IPSPs) relative to WT, leading to extended activation period (EPSP-to-IPSP time). **<u>All panels:</u>** nWT=11, nR6/2=10, mice. Asterisks indicate results from two-tailed t-test; *=p<0.05; **=*p*<0.01; ***=*p*<0.001; ****=p<0.0001; *ns*=non-significant. Hash signs indicate results from two-way ANOVA test, ^#^=p<0.05, ^####^=p<0.0001. All Data is shown as average ± SEM.

### Patterns of human HD pathology align with results in mice

Our findings in R6/2 mice revealed a striking vessel-dependent pattern of astrocyte activation, but a key question remained: does HD pathology in humans exhibit the same vessel- and region-specific heterogeneity? We analyzed post-mortem brain tissue from HD (n=13) and non-HD patient donors (n=7) (Supplementary Table 3). To avoid confounds such as meningeal damage or subarachnoid hemorrhage, we selected cortical sections with well-preserved vessels and meninges. Since murine brains lack prominent white matter, we focused our analysis on grey matter, ensuring better comparability with R6/2 data. Similarly to R6/2 mice, human HD brains exhibited vessel-specific astrocyte activation. Using Collagen IV to mark blood vessels, α-SMA for arterioles, and GFAP for astrocytes, we found that astrocyte activation was strongest around pial and penetrating arterioles, forming dense halos of hypertrophic astrocytes (Fig. 6A, B). In contrast, venules showed minimal astrocyte activation (Fig. 6C-D, Extended Data Fig. 7A-B).

**Figure 6.**
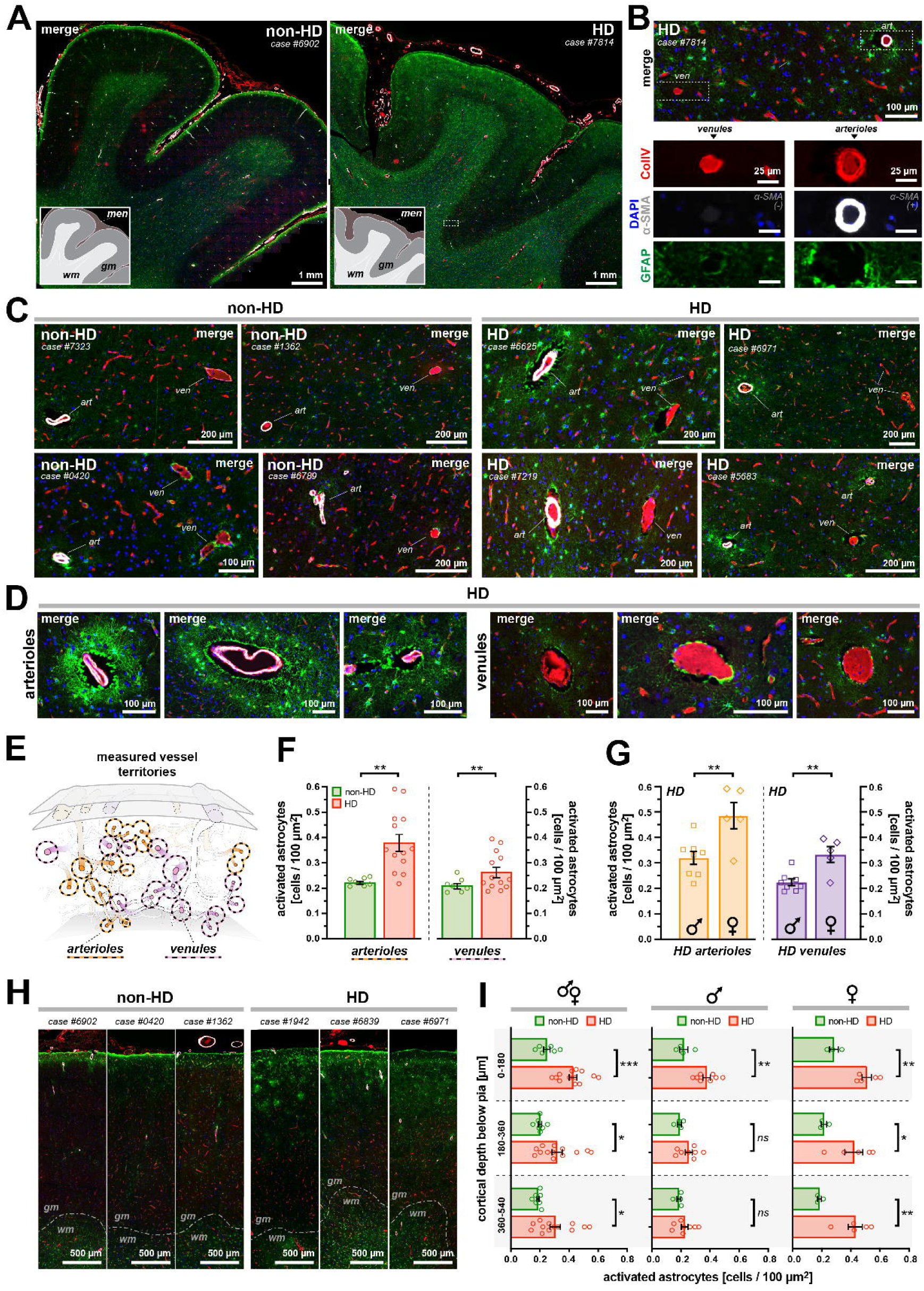
HD patients exhibit arteriolar susceptibility, matching vascular pattern of astrocyte and BBB pathology in mice. (A-B) Representative immunostained cortical sections from non-HD and HD patients. Collagen IV marks vasculature (red), DAPI labels nuclei (blue), α-SMC selectively identifies arterioles (white), and GFAP stains activated astrocytes (green). Small insets in (A) highlight anatomical features of the displayed regions. **(C)** Images from individual HD patient cases showing neighbouring arterioles and venules. HD-affected brains show pronounced astrocyte activation near arterioles, with less affected venules. This arterioral enrichment pattern was absent in non-HD brains. **(D)** Magnified images of arterioles and venules from HD cases showing the extent of astrocytic activation at arterioles. **(E)** Schematic drawing of human cortical brain vasculature depicting the location and spatial orientation of analysed arteriolar and venular territories. **(F)** HD disease phenotype is associated with the enrichment of activated astrocytes at arterioles, with a moderate increase in proximity to venules. **(G)** HD vascular phenotype is sex-driven, with females being significantly more affected than males at both arterioles and venules. **(H)** Images from individual HD patient cases showing enrichment of activated astrocytes at the apical layers of the cortex gray matter. The dashed line shows the anatomical boundary between the gray matter (*gm*) and white matter (*wm*). **(I)** Astrocyte activation is most pronounced in superficial cortical layers, decreasing with depth. This effect is sex-dependent, where females show activation across all analyzed depths and a widespread and severe pathology. **<u>All panels:</u>** *men* = meninges; *gm* = gray matter; *wm* = white matter; nnon-HD= 7; nHD= 13 cases. Data is shown as average ± SEM; two-tailed t-test; *=p<0.05; **=*p*<0.01; ***=*p*<0.001; *ns*=non-significant.

While the number of penetrating and ascending vessels was fewer due to anatomical differences, the overall pattern resembled that observed in R6/2 mice (Extended Data Fig. 6C). To quantify this, we measured astrocyte density within a standardized 20-μm radius (Fig 6E, Extended Data Fig. 8A-B)^25^. The most pronounced astrocyte activation occurred around arterioles, whereas venules were only moderately affected (Fig. 6F). Interestingly, female HD patients exhibited ∼40% greater astrocyte activation than males, an effect observed around both arterioles and venules (Fig. 6G). This sex-driven effect was absent in non-HD brains, where astrocyte densities were comparable between males and females, regardless of vessel type (Extended Data Fig. 8C). Notably, activation levels did not correlate with CAG repeat length or Vonsattel HD severity grade, suggesting that astrocyte activation is a fundamental, rather than stage-dependent, feature of HD (Extended Data Fig. 8D). Internal controls confirmed that age at death and post-mortem interval were not confounding factors in the observed GFAP+ changes (Extended Data Fig. 8E,F).

Next, we examined whether human astrocyte activation follows a gradient, as observed in BCSFB/BBB vulnerability in R6/2 mice, where superficial cortical layers exhibit greater pathology than deeper layers. Given that human cortical gray matter is ∼3× thicker than in mice (2.5-3.0 mm vs. 1 mm)^38,39^, we adjusted our stratification from 60 μm increments in mice to three proportional layers in humans (0-180 μm, 180-360 μm, 360-540 μm) (Extended Data Fig. 8G). Our results revealed that, as observed in R6/2 mice, astrocyte activation in human HD brains was strongest in superficial cortical layers of gray matter and diminished with depth (Fig. 6H-I). Notably, this effect was also sex-dependent. In males, significant astrocyte activation was confined to the superficial 0-180 μm layer, whereas in females, activation extended across all analyzed depths, indicating broader and more severe pathology (Fig. 6I). Thus, HD pathology in humans follows a conserved vessel- and region-specific pattern, where arterioles drive astrocyte activation while venules remain relatively spared, with strong sex-specific vulnerabilities.

## DISCUSSION

In the healthy brain, endothelial tight junctions (TJs) restrict paracellular permeability, preventing systemic macromolecules from entering the parenchyma. Disruption of this function accompanies stroke^32^, epilepsy^40^, and sepsis^41^, and chronic disorders such as Alzheimer’s and Parkinson’s disease (AD, PD)^42–47^. Recent work also implicates age-related microvascular alterations as a contributing factor in BBB dysfunction, reinforcing its role as both a target and a mediator of CNS pathology^10^. However, unlike other proteinopathies, BBB and BCSFB dysfunction in HD has received so far little attention. The few existing studies report conflicting results, with some identifying increased permeability^16,48^, while others find no leakage, even in the late stage of the disease^19,49^. Our current findings resolve these contradictions, We here show that vascular barrier dysfunction in HD is not a global phenomenon but rather varies by barrier and vessel type (summarized in Extended Data Fig. 9). Despite widespread mHtt expression across all vessel segments and barrier types^16^, paracellular leakage appeared selective to pial vessels forming BCSFB, whereas BBB at capillaries remained functionally intact. This vulnerability may be attributed to structural differences, as capillary BECs possess stronger TJs and anti-inflammatory defenses compared to larger vessels^1,2^. Moreover, when ultrastructural TJ alterations occur in capillaries, they do not always translate to increased permeability, as evidenced in iPSC models of juvenile HD^50^ and other CNS pathologies^51^. By contrast, large pial vessels – arterioles, are particularly susceptible to mechanical stressors like shear forces, exacerbating vascular dysfunction^52^. At the same time, venules, with inherently weaker intercellular junctions, are structurally predisposed to inflammation-induced permeability and have heightened responsiveness to inflammatory cytokines, further destabilizing their barrier integrity^53–55^.

The restricted paracellular permeability and low level of vesicular transport is a key function of the BBB and BCSFB ^10,29,32,51,56^. It is only recently that the varying levels of AMT across different vessel types under physiological and pathological conditions were recognized, but the significance of the AMT is not clear^6,7,10^. In a healthy brain, the AMT is strongly suppressed by Mfsd2a (major facilitator superfamily domain containing 2a), the key regulator of Caveolin-1-dependent transport vesicle formation^30^. Impairment of the Mfsd2a pathway leads to an increased turnover rate and amount of transport vesicles, while the vascular networks appear normal, and TJs are preserved^30^. The second key regulator is S1PR1 (sphingosine-1-phosphate receptor-1), located at the luminal side of BECs^6^. The ongoing homeostatic activity of S1PR1 maintains low AMT, especially in arterioles^6^. Here, we observed disinhibition of AMT in R6/2 mice, with arterioles showing the highest susceptibility. This dataset converges with earlier reports on increased transport in post-mortem HD brains^16,48^ and clearly defines the vascular hot-spots, route, and the mechanism of entry. Notably, subpial arterioles exhibited an inverse relationship between paracellular permeability and AMT, with high permeability correlating with modest AMT elevation and vice versa. This highlights the need to consider both paracellular and transcellular mechanisms in HD vascular pathology. The importance of this is evident, as regardless of the transport route, a compromised paracellular barrier and disinhibited AMT both increase the exposure of the CNS to pathological molecules.

Earlier studies reported the protective effects of S1PR1 agonists during inflammatory conditions^6,57^, including HD^58^, with fingolimod being a clinically approved S1PR1 agonist to treat ALS^28^. We observed that selective S1PR1 agonism significantly reduced BBB leakage, albeit the degree of rescue effect depended on the vessel type. The treatment was most effective in the most affected vessel types, both for paracellular leakage (pial vessels), and for disinhibition of AMT (arterioles). This response may be linked to the preferential expression of S1PR1 in arterioles^3^, and much stronger reliance on S1PR1 signaling for maintenance of the BBB in arterioles than in capillaries^6^. In turn, the lack of rescue at capillaries may be explained by a higher dependence on Mfsd2a signaling pathway to regulate the AMT^30^. Nonetheless, the ability of S1PR1 agonists to reduce paracellular leakage and AMT increase in sub-pial vessels emphasizes their potential in treating vascular dysfunction in HD^16–20^.

Blood/CSF proteome fingerprinting provided important insight into the identity of cells affected by the HD. Although MS alone cannot directly map vessel types, correlating protein biomarkers with known expression patterns and vessel-specific functions allowed us to infer distinct vascular mechanisms in HD barrier pathology. MS highlighted immune infiltration and complement-driven endothelial injury, primarily at venules, alongside the active involvement of AMT, particularly in arterioles, which reflects vesicle-driven transport defects. Treatment with the S1PR1 agonist partially normalized this proteomic pattern, corroborating imaging findings of spatially segregated junctional disruptions and vesicular overactivity. Further studies may utilize this information to design precise, barrier-type, and vessel-type targeted therapeutic interventions.

The invasion of blood-derived factors into the brain microenvironment triggers astroglial activation. Notably, albumin activates astrocytes, promoting via TGF-β signaling a senescent, pro-inflammatory phenotype^59^. Vascular and glial dysfunctions emerge as early contributors to HD pathogenesis, detected even in presymptomatic HD carriers (Grade 0 HD), where striatal astrocytes exhibit increased GFAP expression, indicative of early activation^60^. Here, astrocyte reactivity was increased locally at regions exhibiting BCSFB leakage. Although activated astrocytes may locally release neurotrophic factors and antioxidants, they can also acquire neurotoxic properties, secreting reactive oxygen species and excitotoxic metabolites such as quinolinic acid, driving cell damage^61^. Additionally, chronic astrogliosis impairs essential astrocytic functions, including glutamate and K+ uptake, promoting synaptic hyperexcitability and excitotoxicity^61^. In HD, the cerebral cortex exhibits hyperexcitability, even at the early stages of the disease^62^, with increased sensitivity to glutamatergic excitotoxicity during disease progression^63^. Consistent with this, we observed that R6/2 mice exhibited increased neuronal excitability, a faster onset of EPSPs, and delayed IPSPs, indicating persistent neurological deficits. This may be linked with activated astrocytes, which in reactive states exhibit impaired neurotransmitter clearance, leading to prolonged synaptic dwell time of neurotransmitters and disrupted ionic homeostasis due to dysregulated Kir4.1 channels and gap-junction uncoupling^61^.

In summary, this study delineates a zonation of vascular dysfunction in HD, with a promising therapeutic avenue. Our molecular, structural, and functional data provide a framework for targeted interventions, while the identification of vessel-type-selective proteomic markers offers a path toward personalized diagnostics. Further studies are warranted to expand on these findings and translate them into human therapeutics, both in HD and, by extension, other neurodegenerative disorders.

## Supporting information

Supplementary Videos Legends, etc.

## ACKNOWLEDGMENTS

Micael Lønstrup for his excellent assistance with animal surgeries and Kjeld Møllgård for his insightful comments on the manuscript. This study was supported by the Læge Sofus Carl Emil Friis og Hustru Olga Doris Friis’ Legat, Lundbeck Foundation (#R392-2018-2266 and #R345-2020-1440), the Danish Medical Research Council (#1030-00374A), the Independent Research Fund Denmark, and the Novo Nordisk Foundation (#117272 and NNF14CC0001).

## ONLINE METHODS

### Ethical considerations

Formalin-fixed paraffin-embedded 5µm-thick cortex sections from Huntington’s disease patients and age- and sex-matched controls were acquired through the University of Washington Biorepository and Integrated Neuropathology Lab (PI: C. Dirk Keene, MD, PhD) under the REC of the CHU de Québec ethical approval A13-02-1138. All animal surgical and imaging protocols were approved by The Danish National Committee on Health Research Ethics. The experiments followed ARRIVE guidelines and conformed to the standards established by the European Council Convention for the Protection of Vertebrate Animals Used for Experimental and Other Scientific Purposes (permit #2019-15-0201-01655).

### Animal Model

We used 23-24-week-old R6/2 mice corresponding to the late stage of the disease, with their age-matched WT littermates as controls. The R6/2 mice colony exhibited onset of the disease at ∼16 weeks of age (266 to 393 CAG repeats), resulting in a slower disease progression compared to other R6/2 mice with shorter, e.g., 150 CAG repeats^64^. The mice were genotyped for the mutant huntingtin gene (mHTT) as previously described^65^. All animals underwent microsurgical preparation for imaging, i.e., tracheotomy and catheterization of femoral vessels to monitor and maintain the physiological state of the BBB, and craniotomy for 2PM imaging and the placement of EcoG electrode (Fig. 1A). The *in vivo* experiments were performed to obtain from each animal a readout of paracellular permeability, adsorptive-mediated transcytosis, astrocyte count, and measurements of both cerebral and systemic electrophysiology. All animals were investigated in a strict age regime, with the age differences between animals no larger than two weeks, and age-matched healthy littermates (B6CBA mice, referred to as WT) used as controls. The animals were housed in climate-controlled ventilated cages (at 50 ± 10% relative humidity, at room temperature), with 12 h light/ 12 h dark cycle with standard (constant, low) noise level and *ad libitum* access to food and water. The animal housing facility has been accredited by the Association for Assessment and Accreditation of Laboratory Animal Care (AAALAC), and the Federation of Laboratory Animal Science Associations (FELASA).

### Surgical preparations

The animals underwent microsurgical preparation, as described before, with minor modifications^8^. Briefly, the animals were initially sedated via intraperitoneal (i.p.) injections of xylazine (10 μg/g_animal_) and ketamine (60 μg/g_animal_). Subsequently, and through all steps of surgery, the anesthesia was maintained with recurrent i.p. injections of ketamine (30 μg/g_animal_) administered at 20–25 min intervals. A tracheotomy was performed to facilitate mechanical respiration with a stroke volume of 180–220 μL at the frequency 190–240 strokes /min (MiniVent Type 845 respirator, Harvard Apparatus). The inhaled air was supplemented with oxygen at the rate of 1.5–2 mL/min. Next, dual catheterization was performed. The first catheter was inserted into the left femoral artery to administer substances and monitor the mean arterial blood pressure (MABP; BP-1 Pressure Monitor, World Precision Instruments). The second catheter was inserted into the femoral vein to sustain anesthesia during imaging. Post-catheterization, all wounds were surgically closed, and the animal was turned to the prone position for a craniotomy. The scalp was removed, and the periosteum was cleared with a cotton swab soaked in FeCl_3_ solution. Next, the cranium was attached to a custom metal head plate using glue (Loctite Adhesives). A craniotomy was performed above the right somatosensory cortex, at the coordinates 3 mm laterally and 0.5 mm posterior to the bregma, with Ø = 4 mm, using a diamond dental drill at 4500 rpm. The bone flap was carefully elevated, the dura mater excised, and a low-melting point agarose (type III-A, Sigma-Aldrich) 1% solution in artificial cerebrospinal fluid (aCSF; in mM: NaCl 120; KCl 2.8; Na_2_HPO_4_ 1; MgCl_2_ 0.876; NaHCO_3_ 22; CaCl_2_ 1.45; glucose 2.55; pH = 7.4) was applied to the cerebral surface. Subsequently, a ∼4 mm square and 0.08 mm thickness imaging coverslip (Menzel-Gläser, 24×66mm, #1.5) was placed to seal the craniotomy, leaving a margin of ∼0.5 mm for the insertion of glass microelectrodes into the cortex. Next, the animal was relocated to the two-photon microscope imaging stage, and anesthesia was switched to a continuous intravenous (i.v.) infusion with α-chloralose (50 mg/kg_animal_/h)(Fig. 1A).

The animal was allowed to rest for ∼25 minutes before imaging to enable the washout of ketamine from the system, which, in contrast to α-chloralose, hinders NMDA receptor-dependent excitatory neurotransmission. To maintain the animal’s physiological state, during all surgical and imaging procedures, the end-tidal CO_2_ (Capnograph Type 340, Harvard Apparatus) and mean MABP were monitored in real-time, and normothermia at 37 °C was ensured via a rectal thermistor-regulated heating pad. Before imaging, a 50 μL blood sample was collected via arterial catheter to assess blood gases (ABL Radiometer), and the respiration volume and rate were adjusted if necessary.

### Drug treatment

The animals were acutely treated with i.p. injections of SEW2871, the sphingosine-1-phosphate (S1P) receptor (S1PR1) agonist. The treatment started five days prior to imaging procedures, and SEW2871 was administered in 24-hour intervals at the dose of 10 μg/g BW in PBS with 10% DMSO. The WT and control R6/2 group received the corresponding injections with DMSO solution only. The total number of injections was six per animal, with the final injection performed on the experimental day, three hours before the imaging procedures.

### Two-photon *in vivo* imaging setup

The two-photon (2PM) *in vivo* imaging was performed using SP5 upright laser scanning microscope (Leica Microsystems) integrated with MaiTai Ti:Sapphire tunable infrared laser (Spectra-Physics). The images were acquired using an HCX APO L 20 × 1.0 NA water immersion objective. The estimated planar optical resolution was ∼0.53 μm (XY), and the depth resolution was ∼2.5 μm (all estimates for λ_excitation_ = 870 nm). The fluorescence signal was first split by FITC/TRITC dichroic mirror and collected after 525–560 nm / 560–625 nm bandpass filters by two separate NDD multi-alkali photomultipliers (Leica Microsystems). The images were recorded using LAS AF v. 2.4 software (Leica Microsystems) in 16-bit color depth and exported to ImageJ for analysis (v. 1.52a; NIH). 3D reconstructions were performed using volume rendering in Amira v. 6 (FEI Visualization Sciences Group).

### Imaging BBB paracellular permeability

The paracellular permeability assays were performed as described before^6,24^ with minor modifications. Briefly, each animal was bolus-injected with 1% sodium fluorescein (NaFluo; 0.376 kDa; Sigma-Aldrich) at the dosage of 40 µL/20 g_animal_ via an arterial catheter. A hyperstack (Z-stack over time; 4D imaging) was recorded at λ=920 nm excitation for t = 30 min, in bi-directional mode, encompassing the cortical volumes of 775.76 µm x 775.76 µm x 190 µm (XY pixel resolution = 1024 x 1024, Z depth = 38 planes, Z step = 5 µm), with 1 min interval between consecutive Z-stacks.

### Quantifying BBB paracellular permeability

First, to correct for the tissue micromovement during the recording, the timelapse images were stabilized. The hyperstack data has been exported to ImageJ and underwent dimensionality reduction by maximum intensity projection along the Z (depth) axis. The resulting timelapse image has been processed using the *Image Stabilizer* plugin^66^, with *Template Update Coefficient = 0.9; Maximum Iterations = 20; and Error Tolerance 0.0000001*, to obtain the XY transformation coefficients (vector of tissue movement).

Next, using stabilized maximum projection as an anatomical reference, 100 small circular regions of interest (ROIs) were placed uniformly across the brain parenchyma (pROIs) where the signal from vessels was absent both in XY and along the Z axis (Extended Data Fig. 1A). The pROIs’ positions were saved for subsequent use. Then, 10 circular ROIs were placed at the lumen of different pial vessel segments (vROIs) to obtain the average readout of the intensity of fluorophore circulating in the bloodstream over time (Extended Data Fig. 1B).

To obtain the readout from brain parenchyma with respect to the vascular niche, the original hyperstack has been divided into three separate volumes, i.e., superficial *sub-pial* volume (0-60 µm under pia) containing pial and penetrating/ascending vessels; *intermediate* volume (60-120 µm under pia) containing all types of the brain microvasculature, and *capillary bed* volume (120-180 µm under pia) containing primarily the capillaries. Each of these volumes underwent average intensity projection in Z, and for each animal, we analyzed the average projection of *sub-pial* and *capillary bed* volume over time.

Next, we applied previously obtained XY transformation coefficients to stabilize each volume equally. Subsequently, we extracted the baseline fluorescence (at t=0 min) and obtained the readout of fluorescence increase over time. The data was collected from previously placed pROIs (Extended Data Fig. 1A), and then averaged within each animal. To estimate the BBB permeability, the NaFluo signal from each respective volume of parenchyma was divided at each timepoint of the recording by area under the curve (AUC) of the signal obtained from blood vessels (vROIs placed previously on the vessel lumen) (Extended Data Fig. 1B).

### Imaging adsorptive-mediated transcytosis

We assessed the degree of adsorptive-mediated transcytosis (AMT) by imaging the endothelial uptake of the bovine serum albumin (BSA) conjugated to Alexa Fluor488 fluorescent dye (BSA-Alexa488; Invitrogen; emission in green). BSA-Alexa488 was administered i.v. as a single bolus injection (1% in saline; 50 μL /g animal) via a femoral arterial catheter. Immediately after, a hyperstack (Z-stack over time; 4D imaging) was recorded at λ=920 nm excitation, in bi-directional mode and with triple frame averaging, for t = 120 min, encompassing the cortical volumes of 387.5 µm x 387.5 µm x 138µm (XY pixel resolution = 2048 x 2048, Z depth = 55 planes, Z step = 2.52 µm), and with 7.5 min interval between consecutive Z-stacks. Over time, the BSA-Alexa488 appeared at the BBB interface in the form of bright fluorescent puncta, which corresponded to its vesicular uptake (endocytosis). Following that (at t = 135 min), we injected 65 kDa tetramethylrhodamine isothiocyanate-dextran (1% in saline; TRIT-dx; Sigma-Aldrich; emission in red) to delineate a vessel lumen and better distinguish between BSA-Alexa488 in circulation (overlapping with TRITC-dx) and BSA-Alexa488 puncta (Fig. 1A). Lack of TRITC-dx extravasation to the tissue also served as an internal control for the physical integrity of the BBB following the cranial window microsurgery and imaging procedures.

### Quantifying adsorptive-mediated transcytosis

AMT imaging hyperstack data has been exported to ImageJ and an anatomical map has been performed, where all individual vessels segments in the field of view were indexed to belong to one of the following vessel types: pial arterioles (*pA*); penetrating arterioles (*penA*); capillaries *(cap)* post-capillary venules and ascending venules (*pcV/ascV*), pial venules *(pV)* and interconnecting sinuses *(sin)*(Fig. 1B). This anatomical division was based on the vessel location, morphology, branching pattern, diameter, and direction of the blood flow, as previously described^6,8,67^.

At the timepoint 135 post-BSA injection, we performed (i) vessel surface measurements and (ii) AMT puncta count to obtain (iii) AMT puncta surface density, which corresponds to the extent of AMT. Ad (i): each vessel segment was traced in three dimensions using ImageJ *3D measure length* macro (available at http://imagej.net/macros; Jan 2024). The distances were traced manually, always choosing the center of the vessel lumen as an anchor location for the measurement points. Next, for each vessel segment, the vessel lumen diameter was measured at 3-to-5 locations distributed equally along the vessel length (*Line Tool*, ImageJ). The measurements were next averaged to obtain a mean, representative readout. Both, vessel length and diameter were then used to calculate the surface of each respective vessel segment. Ad (ii): the puncta density was manually counted in 3D for each vessel segment in the field of view (*Point* Tool, ImageJ) using the last frame of a recorded hyperstack, i.e., after injecting TRITC-dx into circulation to delineate vessel lumen (at t = 135 min). The selection criteria for a positive identification of the puncta was the presence of the BSA-A488 signal (emission in green) and the absence at the exact location of the signal from other sources (emitting in red), which would indicate that the observed puncta does originate from BSA-A488; but from autofluorescent debris, e.g., lipofuscin in the brain (Fig. 1C). Ad (iii) To calculate AMT puncta density at distinct vessel types, for each mouse, we summed up the puncta count of each respective vessel type and divided it by a sum of their surfaces. This was to eliminate a bias of averaging the density values obtained, e.g., per every single vessel, where one “AMT” - dense vessel with a small surface might skew the results of the whole vessel population. The AMT puncta density has been expressed as an average number of AMT puncta per μm² of the surface of a vessel wall delineated by TRITC-dx.

### Cerebrospinal fluid collection

A 1 ml syringe connected to a polyethylene–tubing–linked glass capillary (outer diameter ≤80 µm) was used for fluid aspiration. Mice were deeply anesthetized with i.p. injection of xylazine (10 μg/g_animal_) and ketamine (60 μg/g_animal_), and mounted onto a stereotaxic frame with a 30° downward head tilt to expose the posterior neck. Following a midline incision and retraction of the overlying muscle, the dura overlying the cisterna magna was gently cleared using sterile PBS and dried with a cotton swab. The capillary was inserted between surface vessels into the cisterna magna. CSF was slowly withdrawn while applying minimal negative pressure to minimize the risk of contamination during collection. Samples were expelled by an air buffer pre-loaded in the syringe and snap-frozen in low-binding Eppendorf tubes on liquid nitrogen and stored at −80 °C for further analyses.

### Mass spectrometry analysis

Samples were stored at −80°C until sample preparation for MS analysis. 5µL of each sample were processed in an SDC buffer (1% w/v sodium deoxycholate, Sigma-Aldrich; 40mM 2-Chloroacetamide, Sigma-Aldrich; 10mM Tris(2-carboxyethyl)phosphine hydrochloride, Sigma-Aldrich; 100mM Tris, Sigma-Aldrich; MS grade water, VWR). Proteins were denatured at 95°C for 10 minutes at 1200rpm. Enzyme was added at 1:100 (w/w), using a master mix of Trypsin (T6567; Sigma-Aldrich) and LysC (125-02543; Fujifilm, WAKO) enzymes, and incubated for 2 hours at 37°C. The reaction was quenched by adding quenching buffer (MS grade isopropanol, Fisher Scientific; 1% w/w trifluoroacetic acid, Sigma-Aldrich) to each sample (pH 2). The sample were transferred to Stagetips containing two layers of SDB-RPS material (Empore) for clean-up. The peptides were washed once with quenching buffer and twice in wash buffer (0.2% TFA, Sigma-Aldrich; MS grade water, VWR) and subsequently dried. The cleaned peptides were eluted using a basic elution buffer (1% ammonia, Merck; 80% MS grade acetonitrile, Merck; MS grade water, VWR) and vacuum-dried to complete dryness at 45°C. Samples were resuspended in a MS buffer (0.1% TFA, Sigma-Aldrich; 5% MS grade acetonitrile, Merck; MS grade water, VWR). Peptide concentration was measured with Nanodrop 2000 and adjusted before injection on the mass spectrometer.

200ng peptides was loaded onto Evotip C18 trap columns (Evosep Biosystems) according to the manufacturer’s instructions. LC–MS/MS analysis was performed on an Orbitrap Astral mass spectrometer (Thermo Fisher Scientific) coupled to an Evosep One system (Evosep Biosystems). Peptides are eluted online from the EvoTips using a 30 SPD (samples per day) method corresponding to a 44-min gradient using a commercial 80mm analytical performance column (Evosep Biosystems). The column temperature is maintained at 55 °C and interfaced with an EASY-Spray Source. For DIA experiments, the Orbitrap Astral mass spectrometer is operated at a full-MS resolution of 240,000 with a full scan range of 380–980 m/z using an automatic gain control (AGC) target of 500% and a maximum injection time of 3 ms. Fragment ion scans (MS/MS scans) are set to 200 windows of 3-Th with a scan range of 150-2000 m/z and a max IT of 5 ms. The isolated ions are fragmented using HCD with 25% Normalized Collision Energy.

Raw files are processed in Spectronaut version 19.5.241126.62635 (Biognosys) in directDIA mode, analyzed against a Mus musculus FASTA file (downloaded from Uniprot on 28-10-2024). Enzyme specificity is set to Trypsin/P, and peptides are filtered to between 7 and 52 amino acids in length with a maximum of 2 missed cleavages. Carbamidomethyl on cysteine is allowed as a fixed modification, and oxidation on methionine and acetyl on the N-terminal as variable modifications. The false discovery rate (FDR) is set at 0.01 on all levels, and cross-run normalization is set to local normalization with row selection by complete identification across all runs (‘Qvalue complete’). Proteomic data processing, including normalization, filtering, statistical analysis, and visualizations, was performed in Python, utilizing an in-house analysis pipeline based on the Clinical Knowledge Graph (CKG)^68^. The data was filtered for at least 70% valid values in at least one group and missing LFQ values were imputed by using a mixed model combining probabilistic minimum imputation and k-nearest neighbor (downshift=1.8 standard deviation and width=0.2, knn_cutoff=0.6, n_neighbors=5)^68^. Protein intensities were transformed to a log2 scale and normalized by the median of each sample.

### Electrophysiology setup, ECoG, and systemic physiology

Brain electrophysiological activity was recorded using a heat-pulled single-barreled borosilicate glass microelectrode (tip Ø = 2–3 μm; inner Ø = 0.86 mm; outer Ø = 1.5 mm; electrode resistance 1.5–2.0 MΩ; Sutter Instrument) with an Ag/AgCl filament, filled with artificial CSF (aCSF). The electrode was inserted 50-70 μm into the cerebral cortex under a glass coverslip, while the reference electrode was placed between the neck skin and muscle (Fig. 4A). The total electrical signal (TES) was filtered with a 0–3000 Hz low-pass filter and amplified 10 x (AP311 analog amplifier; Warner Instruments). The spontaneous brain activity, or alternate current-ECoG (AC-ECoG) component, was further amplified 100 x and filtered with a 0.5 Hz high-pass filter (NL 106 analog amplifier and NL 125/126 analog filter, NeuroLog). Next, the analog data was digitized at 20 kHz using a Power 1401 interface (CED).

Both EcoG and systemic physiology data, i.e., the raw readout from the exhaled CO_2_ (Capnograph Type 340, Harvard Apparatus) and MABP (BP-1 Pressure Monitor, World Precision Instruments), were recorded via Power 1401 interface (CED) in Spike2 software (v. 7.02a; CED).

### Whisker somatosensory stimulation and LFP

The right mouse sensory barrel cortex was activated by thalamocortical stimulation of the contralateral ramus infraorbitalis of the trigeminal nerve, using custom-made, percutaneously inserted bipolar electrodes, as described before^69,70^, with modification. Briefly, the cathode was positioned at the left hiatus infraorbital (i.o.) nerve, with the anode in the masticatory muscles (Fig. 4A). Stimulation was delivered via an ISO-flex stimulator (A.M.P.I.) at an intensity of 1.5 mA for 1 ms in trains of 15 s at frequencies in Hz: 0.5; 1.0; 2.0; 3.0. and 5.0. To minimize the impact of the preceding train of stimulation, the consecutive train stimulation was spaced 1 minute apart. This round of stimulation was repeated thrice for each animal, totaling 15 stimulation trains across five stimulation frequencies per animal. From each single frequency train, we obtained the average local field potential (LFP) readout (discarding the first stimulus impulse). From each LFP readout, we extracted the baseline (50 ms preceding each stimulus artifact spike), the evoked LFP post-synaptic excitatory potential (EPSP), post-synaptic inhibitory potential (IPSP), and their respective latencies (time from the stimulation impulse to the peak of EPSP and IPSP). Next, all values (EPSP, IPSP, EPSP and IPSP latencies) were averaged across three stimulation rounds. The sum of EPSPs and IPSPs for each frequency was calculated by multiplying the average EPSP and IPSP values by the respective stimulation frequency (0.5; 1.0; 2.0; 3.0. or 5.0 Hz) and by the duration of the stimulation train (15 s).

### SR101 astrocyte density measurements *in vivo*

To label the astrocytes *in vivo*, we loaded a heat-pulled single-barreled borosilicate glass micropipette (tip Ø = ∼4 μm; inner Ø = 0.86 mm; outer Ø = 1.5 mm; electrode resistance ∼1.0 MΩ; Sutter Instrument) with 1 mM sulforhodamine 101 dye (SR101; Sigma-Aldrich) in aCSF^34^. The micropipette was inserted into the cortex at a depth of ∼150 µm, and SR101 was extruded in a series of 4-5 intermittent pressure pulses in 4–5 intermittent pulses (2-s pulse; 10-15 psi; pneumatic picopump PV830; World Precision Instruments). After 40 min, the SR101 uptake clearly labeled astrocyte somata and processes. To count the average density of astrocytes for each animal, we performed a series of Z-stacks, encompassing the cortical volume of 387.5 µm x 387.5 µm x 159.7-304.4 µm range (XY pixel resolution = 1024 x 1024, Z depth = 32-61 planes, Z step = 5 µm). The Z-stacks were collected at λ_excitation_=880 nm in a series of three-to-four stacks with increasing laser intensity between consecutive stacks, e.g., 3%; 6%; 12.5%, then 25% of the output power 2.51 W measured at the laser. This was to obtain a high dynamic range of recorded fluorescence signal, which permitted clear identification of astrocytes *in vivo*, especially at the same imaging plane with distinct astrocytes exhibiting varying degree of SR101 uptake. Next, from the total imaged volume, we excluded the volume occupied by pial vessels and the tissue beneath along the whole Z axis (ImageJ). This was because the presence of highly absorbing pial vessels in the tissue might obstruct the detection of astrocytes located beneath vessels in deeper cortical layers (>250 µm). The number of astrocyte somata was manually counted in 3D using Z-stacks collected at different laser power (ImageJ) and expressed as the density of astrocytes / mm^3^ volume of the brain parenchyma.

### Immunohistochemistry in mice

Mice were anesthetized using Xylazine/Ketamine and transcardially perfused at a rate of 10 mL/min with phosphate-buffered saline (PBS) for 1–2 minutes, followed by ice-cold 4% paraformaldehyde (PFA) in PBS for 4–5 minutes. Brains were post-fixed in 4% PFA at 4°C for 24 hours and subsequently cryoprotected in 25% sucrose with 0.1% sodium azide in PBS at 4°C for an additional 24 hours. The brains were then frozen using dry ice and sectioned into 30 μm thick coronal sections using a sliding microtome (Microm HM450, ThermoFisher Scientific).

For immunofluorescence staining, brain sections were initially washed in PBS to remove the antifreeze solution, then preincubated for 1 hour at room temperature (RT) in a blocking solution of 5% normal donkey serum in PBS with 0.3% Triton-X-100. Sections were incubated overnight at RT in PBS with 0.025% Triton-X-100 (PBST) and 3% normal donkey serum containing the following primary antibodies: anti-CD31 (1:400, BD Biosciences, BD550274) to label endothelial cells, anti-α-smooth muscle actin (Anti-Actin, α-Smooth Muscle - Cy3™ antibody, Mouse monoclonal, 1:250, Sigma-Aldrich, A6198) for vascular smooth muscle cells, and anti-glial fibrillary acidic protein (GFAP; 1:500, Agilent Technologies (Dako), Z033429-2) to stain astrocytes. Sections were then washed (3× for 10 min in PBST) and incubated for 1 hour at RT with fluorescent secondary antibodies: Alexa Fluor 488 (1:500, Goat Anti-Rat, Abcam, ab150157), and Alexa Fluor 647 (1:500, Donkey Anti-Rabbit, Abcam, ab150075), all diluted in PBST with 5% normal serum. After another round of washes (3× for 10 min in PBST), sections were stained with DAPI (1:1000, ThermoFisher Scientific) for 10 minutes to visualize cell nuclei, followed by a final washing step in PBS. The stained sections were mounted on gelatin-coated glass slides and coverslipped using an antifading mounting medium (Fluoroshield Mounting Medium with Propidium Iodide, Abcam, ab104129).

### Immunohistochemistry in human brains

Sections were placed on a slide warmer at 65 °C for 20 minutes to remove the paraffin, followed by defatting in Citrisolv (Fisher Scientific, cat# 22143975) and rehydration in a descending bath of ethanol solutions. Sections were then rinsed with MilliQ water before an antigen retrieval was performed with a solution of citrate buffer (sodium citrate tribasic 10mM [MilliporeSigma, cat# W302600], EDTA 2mM [Fisher Scientific, cat# BP-2482], Tween-20 0.05% [Fisher Scientific, cat# BP-337]) pH 6.2 at 80°C from 20 min. After a cooling period of 20 min at room temperature (RT) in citrate buffer, sections were washed with potassium phosphate-buffered saline (KPBS), incubated in peroxide 3% for 30 min, and then immersed in blocking solution (1% bovine serum albumin [BSA, BioShop Canada, cat# ALB001], 0.1% Triton X-100 [MilliporeSigma, cat# T8787], 5% Normal donkey serum [NDS, MilliporeSigma, cat# D9663] in KPBS) for one hour at RT. After additional washes, sections were incubated overnight at 4°C with a mix of primary antibodies in blocking solution; goat anti-collagen IV (1/500; MilliporeSigma, cat# AB769), rabbit anti-GFAP (1/500; Dako, cat# Z0334) and mouse anti-αSMA conjugated with Cy3 (1/200; MilliporeSigma, cat# C6198). The following day, sections were washed with KPBS and incubated with secondary antibodies diluted in blocking solution; donkey anti-goat Alexa Fluor 488 (1/500; Invitrogen, cat# A11055) and donkey anti-rabbit Alexa Fluor 647 (1/500; Invitrogen, cat# A31573) for 2.5 hours at RT. After subsequent washes in KPBS, sections were incubated with DAPI (1/5000, DAPI 5mg/mL, Invitrogen, cat# D3571) for 10 min, washed in KPBS and autofluorescence was quenched using a solution of Sudan Black (MilliporeSigma, cat# 199664) 0.1% in ethanol 70%. Sections were washed in KPBS to remove the excess of Sudan Black, rinsed in MilliQ water and coverslipped with Fluoromount-G (Invitrogen, cat# 00-4958-02). After a drying period, scans of the entire section stained for each case were acquired with an AxioScan.Z1 (Zeiss) using Zen software (ZEN 3.1, Zeiss).

### Statistical analyses

The sample sizes, including the treatment group size, were estimated based on our previous 2PM studies of paracellular permeability, transcytosis, and electrophysiology *in vivo*^6,8,69^. The *in vivo* experiments were performed to obtain from each animal a readout of paracellular permeability, adsorptive-mediated transcytosis, astrocyte count, and measurements of both cerebral and systemic electrophysiology. All statistical analyses were made comparing independent biological replicates, i.e., individual mice. For comparisons between the groups, first, the data was tested for normality using the D’Agostino-Pearson test, and unpaired two-tailed Student’s t-test or two-tailed Mann– Whitney tests were used for normally and non-normally distributed data, respectively. For testing multiple hypotheses, Bonferroni post-hoc correction was used. To test for the difference between more than two means, we used two-factor ANOVA (electrophysiological recording). The statistical analyses were performed using Prism v.10.2 (GraphPad). Data were plotted using Prism v.10 and OriginPro 2018 (OriginLab Corporation). No outlier testing and elimination has been performed to avoid inflation of type I error rates. Proteomic identification of differentially expressed proteins between different groups was executed using t-tests with permutation-based threshold FDR<0.05 and S_0_-value=0.3 at 250 randomizations for each comparison^68,71^. Test results were considered statistically significant for *p* values or corrected *p* values <0.05. The data was presented as average ± standard error of mean (SEM), unless stated otherwise. The imaging experiments could not be performed in a blinded manner due to overt differences between WT and R6/2 phenotypes, but the imaging procedures and data acquisition parameters were the same across all tested groups. The electrophysiology, paracellular permeability, AMT, and cell-count analyses were performed in semi-randomized order (by the ascending ID number of the the animal) by an investigator blinded to experimental conditions. The analyses involving immunohistochemistry were performed in a blinded manner for both R6/2 mice and human brains.

### Model considerations

We chose the R6/2 transgenic mouse model of HD that harbors the human mutant huntingtin gene (mHtt)^12,13,64^, with early onset and rapid progression of disease^12,13^, that exhibits a close resemblance of vascular pathology to changes observed in HD patients, including cell-type distribution of mHtt aggregates^16^, and with similar changes in the transcriptional profile^72^. The R6/2 model, though not replicating all aspects of HD, is well-established for studying pathogenesis and interventions. Its transgene expression, driven by the human huntingtin promoter, achieves ∼75% of endogenous huntingtin levels^65,73^. Importantly, mutant huntingtin (mHtt) is present in all major neurovascular unit (NVU) components, mirroring its deposition in HD patients^15,16^.

### Methodological considerations

The 2PM allowed us to assess changes in microvascular function in the living, intact brain with superior spatial and temporal resolution over whole-brain imaging techniques such as MRI or PET. However, our method also has limitations. First, the diffusion rate of NaFluo in the brain was too fast to identify whether arterioles or venules contributed most to paracellular leakage in the sub-pial region. On that note, it is unlikely that NaFluo leaked at capillaries and diffused along larger surface vessels via perivascular spaces to sub-pial volume. If this were the case, we would observe a concentration gradient with high NaFluo presence at capillaries and low levels in sub-pial volumes, which was not present in the analyzed images. Second, earlier studies using Evans Blue did not account for its non-selective nature. An increase in Evans Blue accumulation within the brain parenchyma may not indicate increased paracellular permeability but could result from increased AMT, as Evans Blue binds to albumin in the bloodstream^74^. Here, we selectively focused on albumin to get insight into AMT. Although albumin is the main protein undergoing AMT, other proteins like IgGs can also utilize this pathway^10^. Despite that, compared to transmission electron microscopy (TEM), which is typically performed on capillaries due to preservation challenges with larger vessels, our method allowed for the first detailed estimation of AMT in large vessels in HD, avoiding the oversights of *ex vivo* studies or cell cultures that lack the brain’s functional and structural heterogeneity.

