## Supplementary Figures for "Vascular dysfunction in Huntington’s disease is located at the blood-CSF barrier and is rescued by sphingosine-1-phosphate receptor agonist"

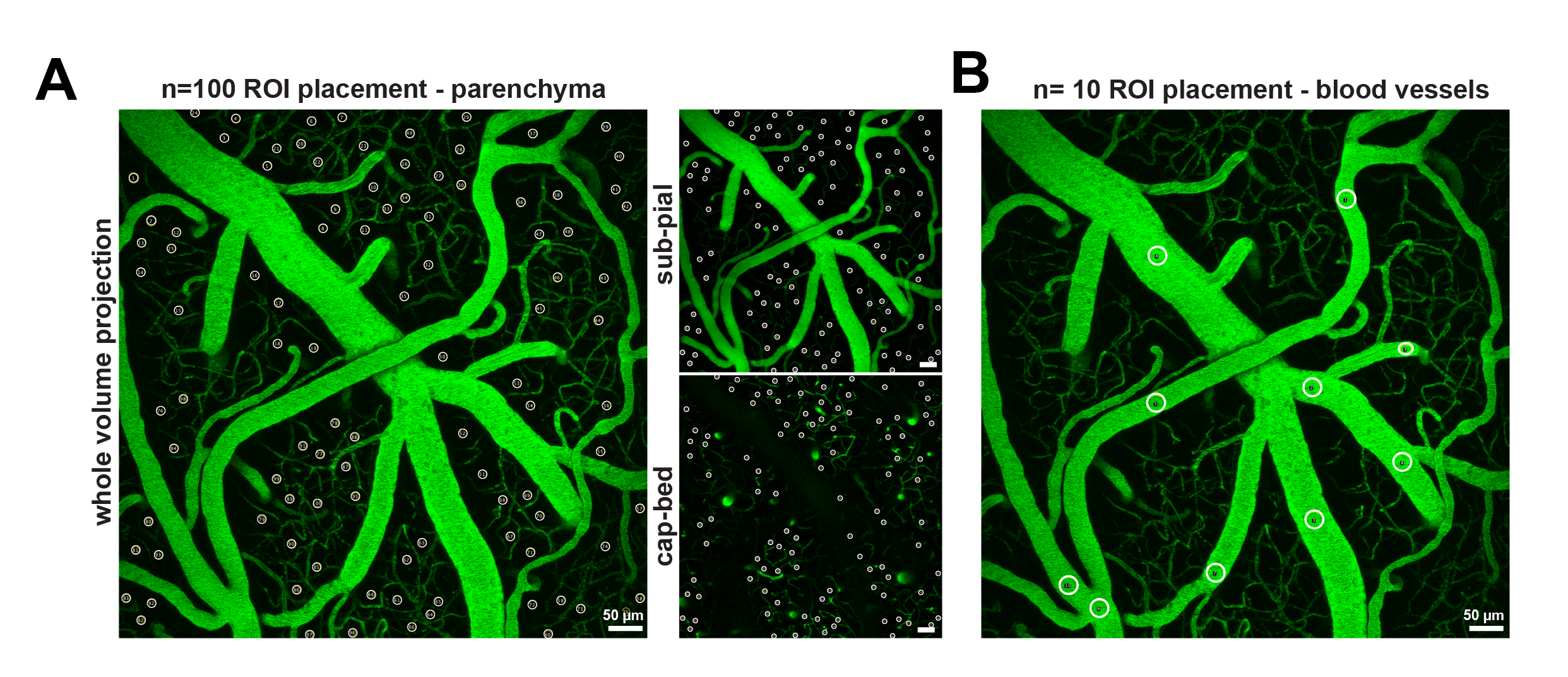

**Extended Data Figure 1. Placement of regions of interest (ROIs) to measure paracellular permeability using 2PM *in vivo*** **(A)** Placement of ROIs in the brain parenchyma (pROIs, white circles) to measure the NaFluo signal increase in the brain. The pROIs are placed in areas devoid of vessels, using as anatomical reference the maximum intensity projection across the whole analyzed volume (left panel). Next, the same set of pROIs is reused for collecting the signal from *sub-pial* and capillary bed *(cap-bed)* volumes of the cortex**. (B)** Placement of the vessel ROIs (vROIs, white circles) to obtain the average signal of NaFluo circulating in the bloodstream. n_pROI_ = 100 / animal, n_vROI_ = 10 / animal; in n_WT_= 9; n_R6/2_= 7; n_R6/2+_= 7 mice.

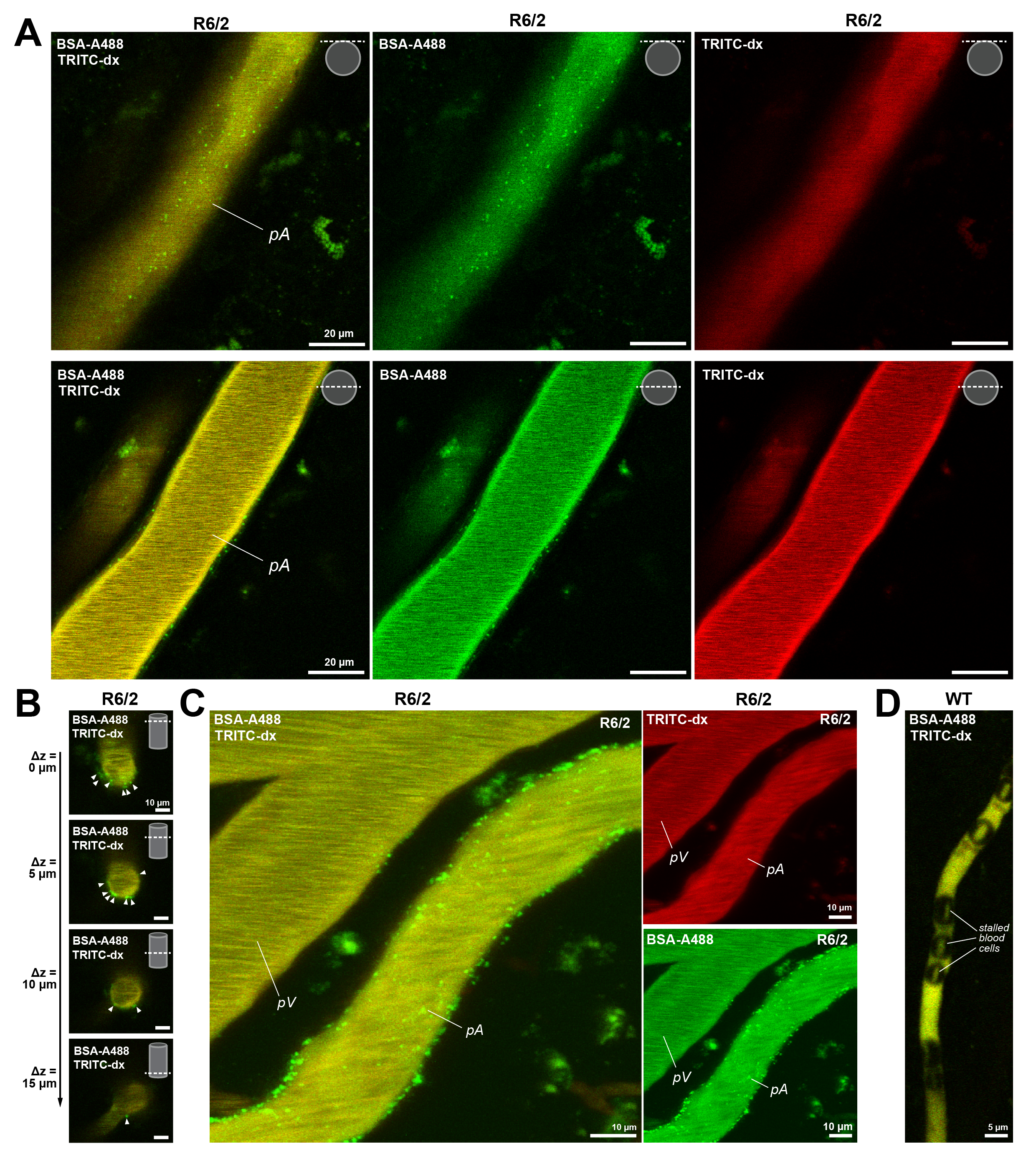

**Extended Data Figure 2. 2PM examples of AMT imaging in vivo**

**(A)** Example images of AMT puncta (BSA-A488, green) imaged in pial arteriole. The upper panels show an arteriole with an imaging focal plane at its pial surface; the lower panels show the same arteriole with the imaging focal plane crossing the broadest section of the vessel. The inserts in the upper-right corner show the location of the focal plane in relation to the vessel cross-section. **(B)** Example images of AMT puncta measured along the penetrating arteriole, with the inset showing the location of the focal plane along the vertically oriented vessel and Δz showing the relative depths of the focal plane. **(C)** Neighboring arteriole and venule show demonstrate contrast in vulnerability (AMT puncta extent) in R6/2 mice. **(D)** Vessels with stalled blood flow do not give rise to AMT increase. The image shows an example of stalled capillary in WT mice, with numerous immobile blood cells obstructing the blood flow. Note the absence of AMT puncta.

**All panels:** *pA* = pial arteriole, *pV* = pial venule.

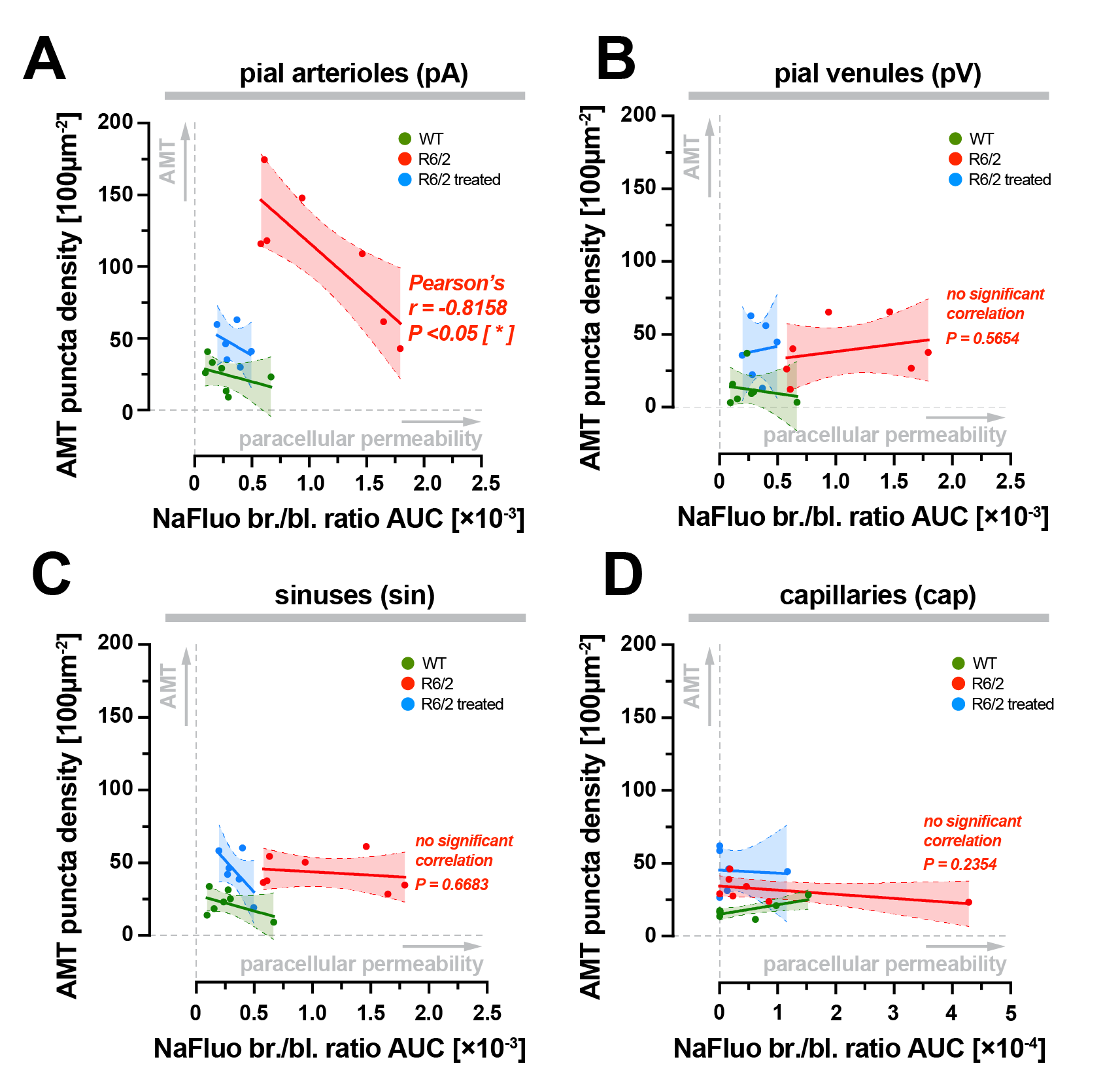

**Extended Data Figure 3. Either AMT or paracellular leakage pathology takes precedence at pial arterioles in R6/2 HD mice. (A-D)** Cross-correlation between the degree of paracellular permeability and AMT in WT, R6/2, and R6/2 treated mice.

**(A)** In pial arterioles (pA), the AMT and paracellular permeability are inversely correlated (Pearson’s r=-0.8168; **p*<0.05), but this relation is not present in other vessel types occupying the same niche **(B-C),** and capillaries **(D)**.

**All panels:** Each point represents a single animal, number of pairs: n_WT_= 7; n_R6/2_= 7; n_R6/2+_= 6 mice; *r* = Person’s correlation coefficient, *p* = the probability that the observed data fits the expected distribution. The thick lines show a linear fit with a shaded area representing a 90% confidence interval. *AMT puncta density* = measure of the extent of AMT; *NaFluo br/bl. ratio AUC* = measure of paracellular permeability, i.e., blood/brain ratio of NaFluo fluorescence increase expressed as area under the curve (see Figure 1H-I). Gray arrows denote the direction of increases in AMT and paracellular permeability.

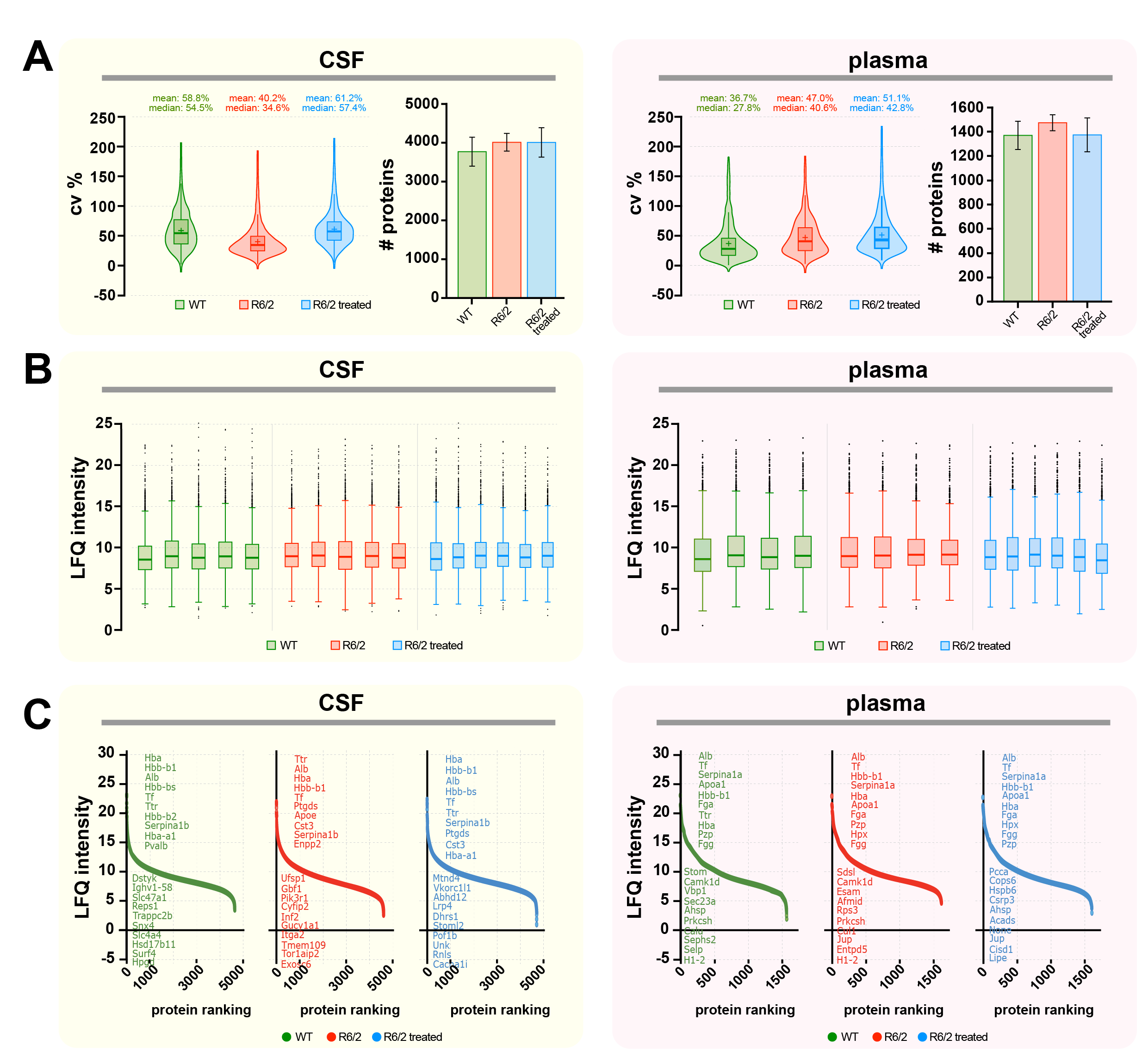

**Extended Data Figure 4. Protein yield and quality control of blood plasma and cerebrospinal fluid samples for mass spectrometry analysis. (A)** The average number of proteins identified and the coefficient of variation (CVs) are illustrated by group for both plasma and CSF. **(B)** Normalized protein intensities across all samples are depicted using box plots for both plasma and CSF to visualize the protein distribution. **(C)** The ranked proteins based on intensity are shown to illustrate the dynamic range and the top 10 highest and lowest expressed proteins by mass spectrometry.

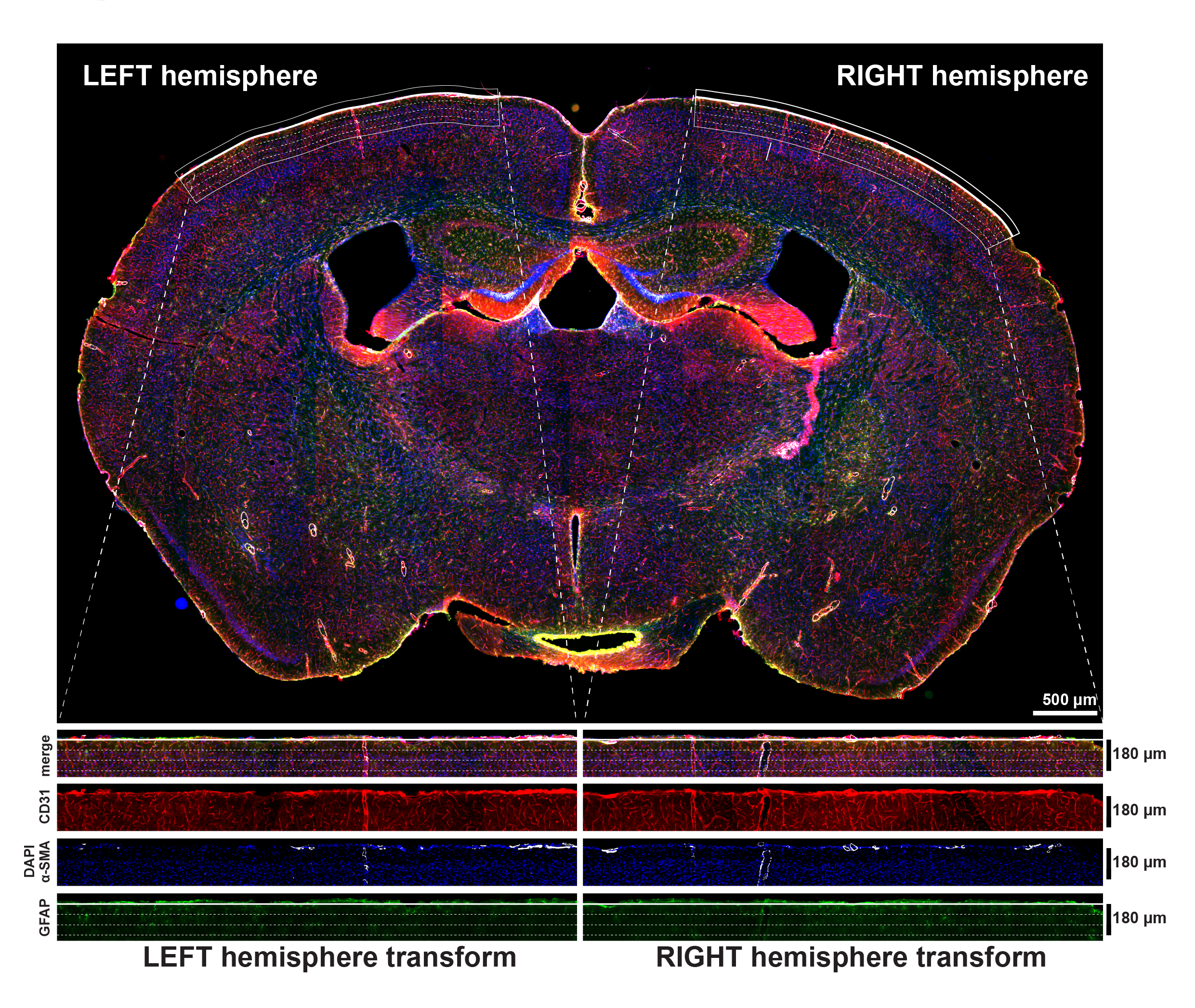

**Extended Data Figure 5. Mouse immunohistochemistry.** Example of a whole-brain immunofluorescence image from the cortical section of an R6/2 mouse. The superficial areas of both hemispheres (a bent ribbon-like structure) underwent non-rigid deformation (straightening) to assess the location of GFAP+ astrocytes in three 60 µm intervals (parenchyma, dashed lines) relative to the brain surface (pia, bold continuous line).

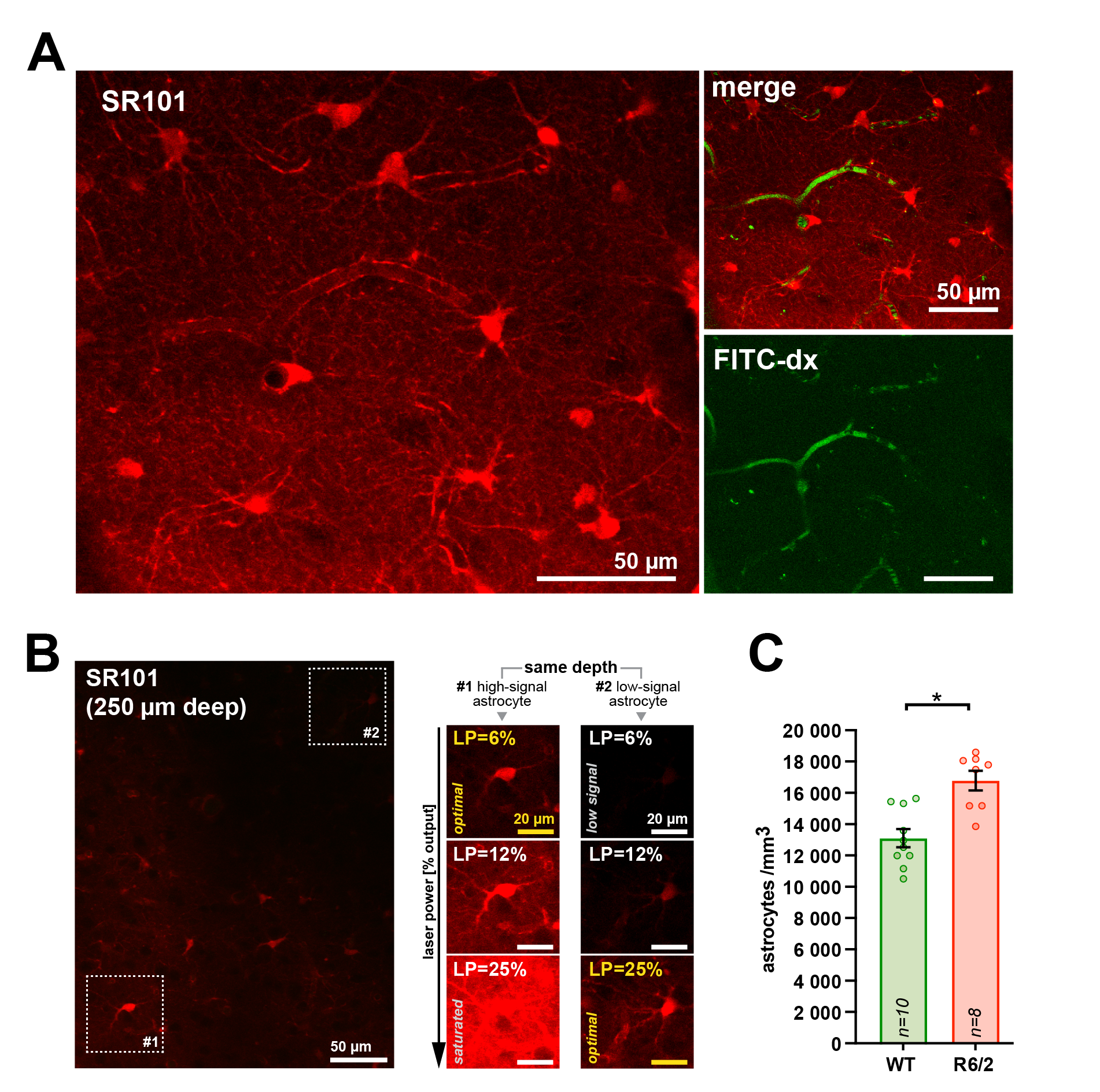

**Extended Data Figure 6. Measurements of astrocyte density *in vivo* (A)** Example images of astrocytes labeled *in vivo* with an intraparenchymal injection of SR101, a pan-astrocytic fluorescent marker. **(B)** Due to the varying extent of SR101 uptake by astrocytes, the images were collected with increasing laser intensity (here: 6-25% of the output power 2.51 W measured at the laser) to obtain the optimal detection range for all, bot high-signal and low-signal astrocytes. **(C)** Compared to WT, both R6/2 and R6/2 treated (R6/2+) mice exhibit increased astrocyte density in cortical parenchyma. n_WT_= 10; n_R6/2_ = 8. Data is shown as average ± SEM; two-tailed t-test; *=p<0.05.

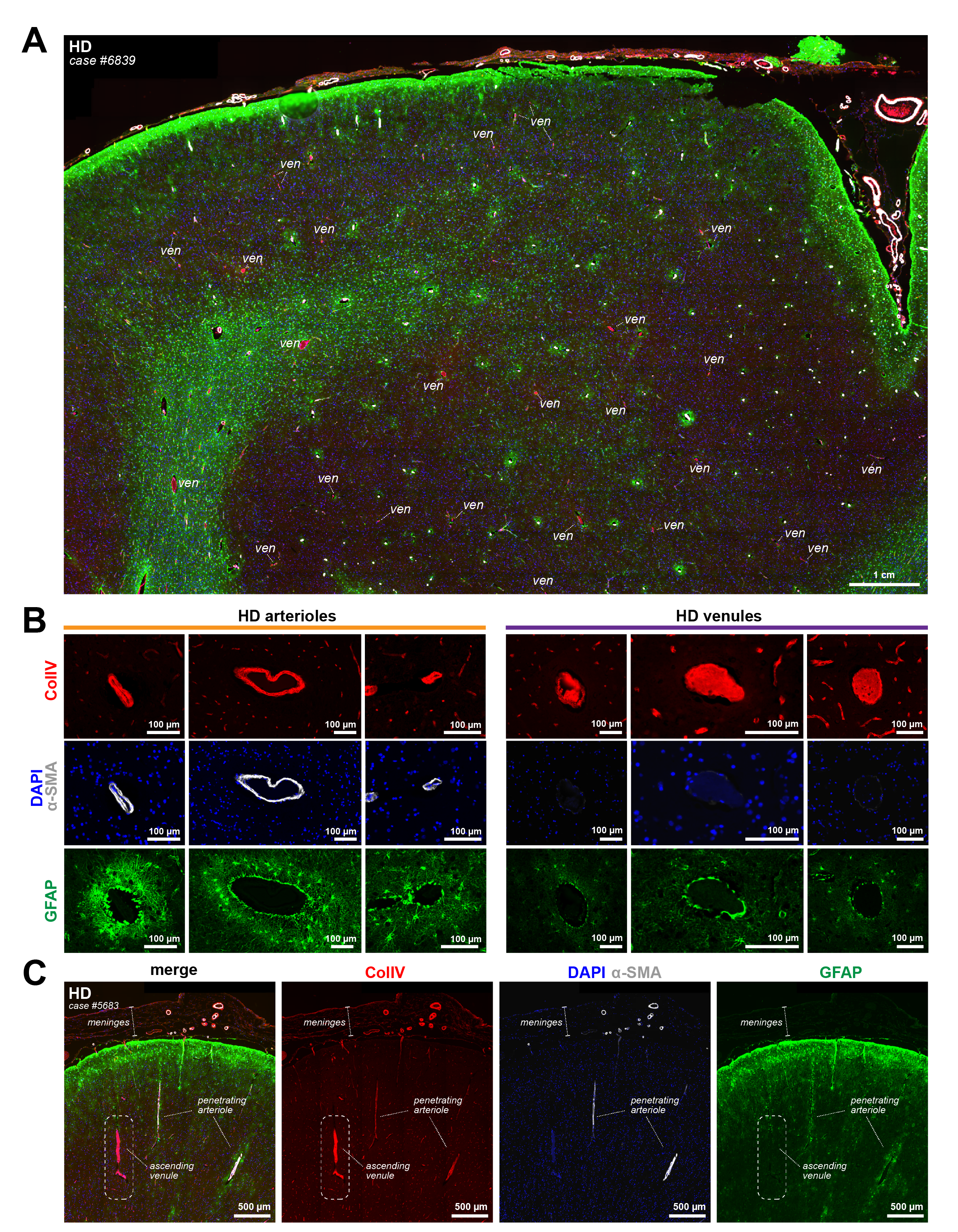

**Extended Data Figure 7. Immunohistochemistry on *post-mortem* human HD brains. (A)** Immunofluorescence image of the cortical region from HD brain depicting a dense network of arterioles and venules within the field of view. The image demonstrates preferential astrocyte enrichment in proximity to arterioles, with minimal association observed in venules. For clarity, only vessels identified as venules are labeled (*ven*), while all unlabeled large vessels correspond to arterioles. **(B)** Representative images from Fig. 4D presented as individual fluorescence channels. GFAP+ astrocytes exhibit a distinct perivascular halo surrounding arterioles, while venules show only minimal astrocytic activation. **(C)** The spatial pattern of astrocyte activation is preserved regardless of vessel orientation. Sparse penetrating arterioles and ascending venules maintain the same trend, with astrocyte enrichment preferentially localized around arterioles.

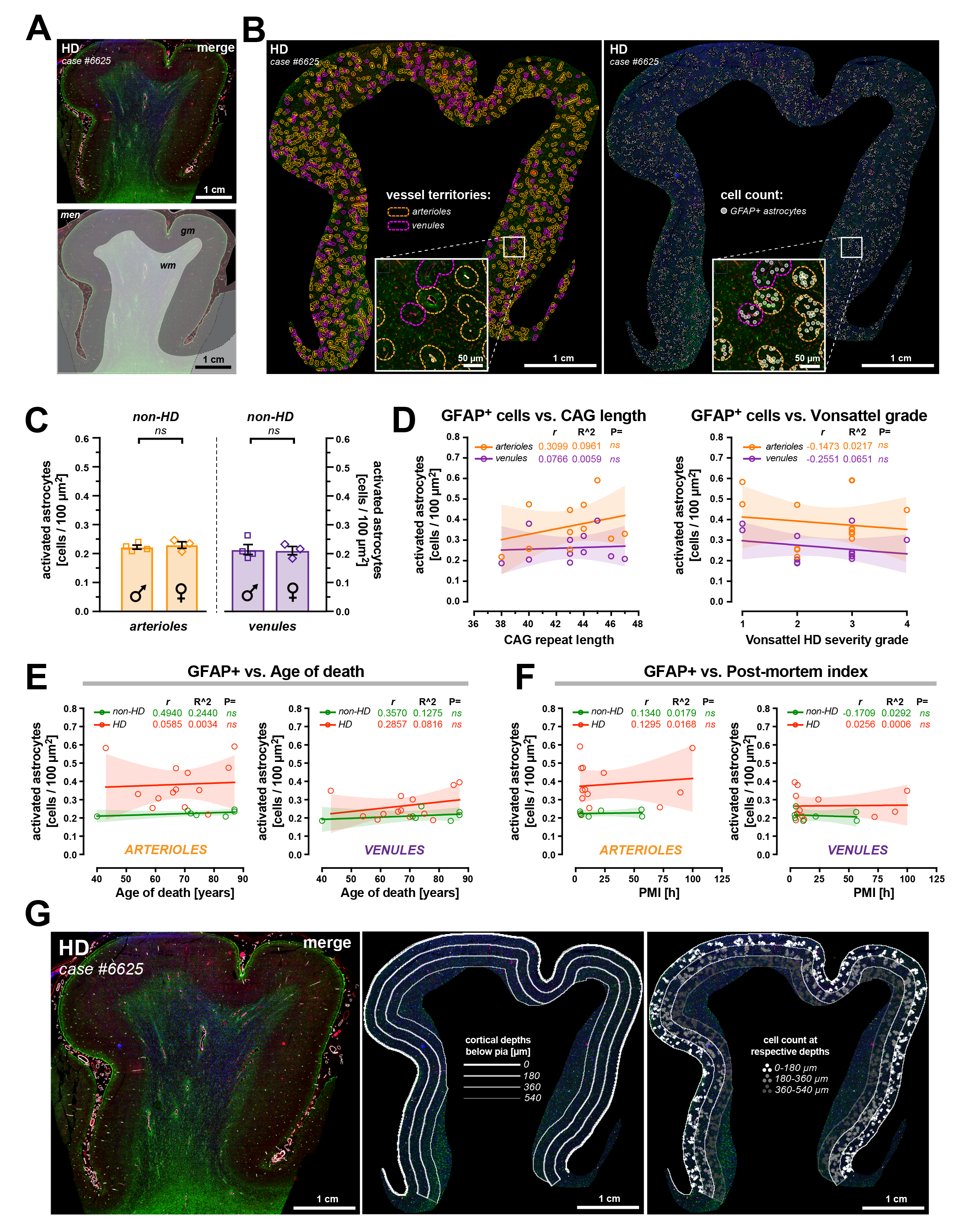

**Extended Data Figure 8. Quantitative analyses of human *post-mortem* brain sections.**

**(A)** Immunofluorescence image of the cortical region from an HD human brain with a schematic representation of anatomical features. **(B)** Gray matter panels corresponding to the brain region shown in (A), delineating arterial (orange) and venular (purple) territories based on a 20 µm cutoff. GFAP+ astrocytes were quantified within each territory, with astrocytes located in overlapping regions excluded from both arteriole and venule counts. **(C)** In non-HD human brains, the density of activated astrocytes is independent of sex and remains comparable between arterioles and venules. **(D)** In HD human brains, the density of activated astrocytes is not influenced by donor CAG repeat length or disease severity. **(E-F)** Internal controls demonstrating that GFAP+ astrocyte density at arterioles and venules is unaffected by the donor’s age at death or post-mortem tissue interval index. **(G)** Panels illustrating the same cortical region as in (A), depicting the analysis of the astrocytic activation gradient in relation to cortical depth.

**All panels:** Each point represents a single donor case, n_non-HD_= 7; n_HD_= 13. The thick lines show a linear fit with a shaded area representing a 90% confidence interval. *r* = Person’s correlation coefficient, R^2^ = goodness of fit. The thick lines show a linear fit with a shaded area representing a 90% confidence interval. Data is shown as average ± SEM; two-tailed t-test; *ns*=non-significant.

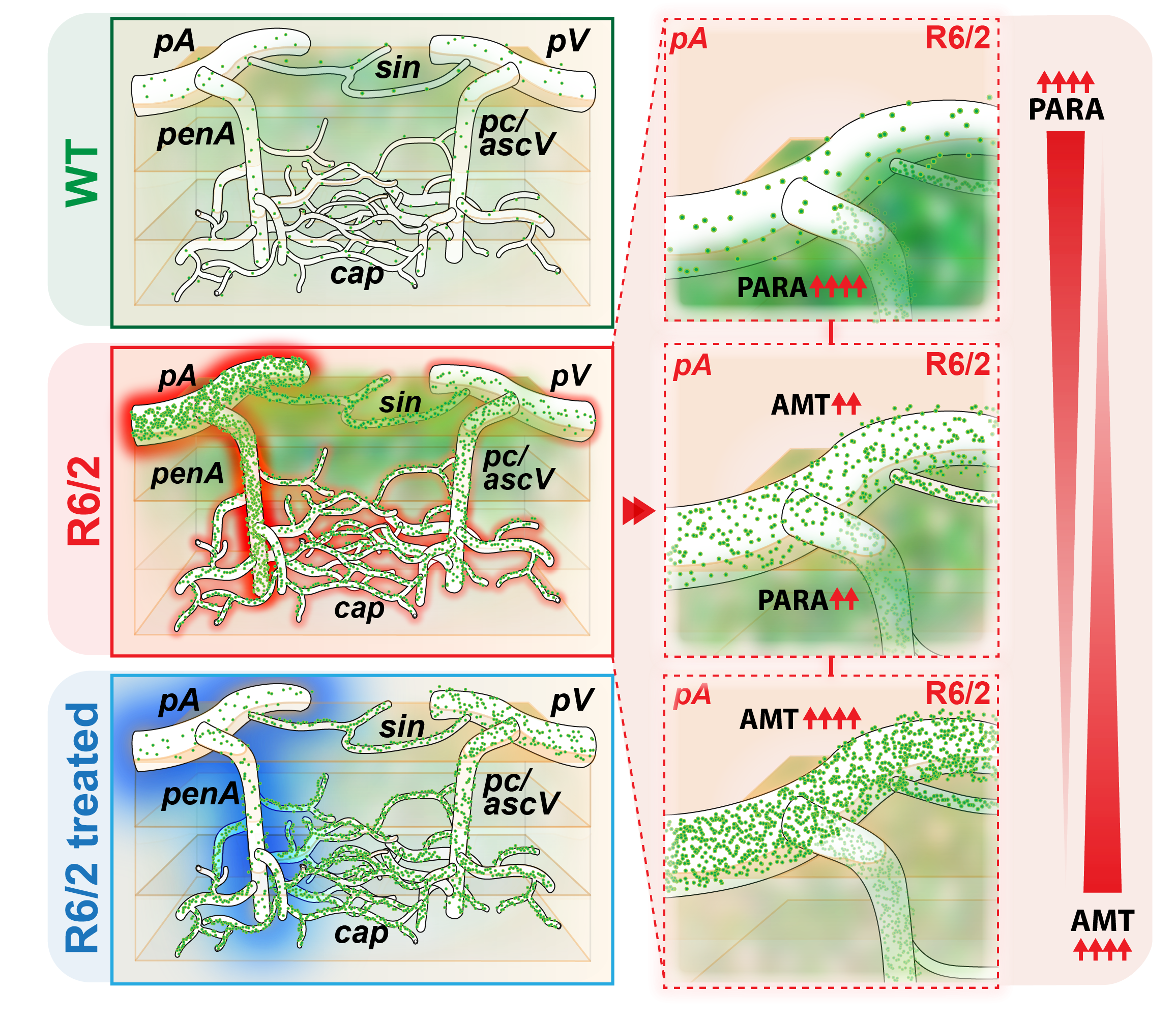

**Extended Data Figure 7. Summary.** HD disrupts brain barriers in a vessel-specific manner. Pial vessels drive paracellular leakage, while arterioles are most susceptible to an increase in transcytosis. S1PR1 agonist restores normal barrier function at pial vessels and reduces transcytosis in arterioles but has only limited effects on other vessel types. Either an increase in paracellular permeability or an increase in AMT is a dominant feature of arteriole dysfunction in HD.
