## Supplementary Videos Legends, etc. for "Vascular dysfunction in Huntington’s disease is located at the blood-CSF barrier and is rescued by sphingosine-1-phosphate receptor agonist"

**SUPPLEMENTARY TABLES**

**Supplementary Table 1:** Summary of 2PM data statistical analysis (attached in .xls file)

**Supplementary Table 2:** Measurements and statistical test summary (attached in .xls file)

**Supplementary Table 3:** Clinicopathological data for patients' post-mortem brain samples (attached in .xls file).

**SUPPLEMENTARY VIDEOS**

**Video 1:** NaFluo paracellular leak into brain parenchyma at distinct cortical volumes, for WT, R6/2, and R6/2 treated mice (R6/2+).

**Video 2:** Motility of AMT puncta on vessels post-injection

**Video 3:** Fibroblast uptake of blood-borne albumin after AMT via the endothelium

**Video 4:** Spontaneous constrictions of large, pial arterioles in R6/2 mice.

**OTHER**

**Mass-spec database for upload**: (attached in .xls file)
